# The penetration ring is a novel infection structure formed by the penetration peg for invading plant cell membrane in rice blast fungus

**DOI:** 10.1101/2024.07.11.603048

**Authors:** Wenqin Fang, Xiaoyu Zai, Jia Chen, Yakubu Saddeeq Abubakar, Qiuqiu Wu, Zhenyu Fang, Xiuwei Huang, Xiang Gan, Daniel J. Ebbole, Zonghua Wang, Wenhui Zheng

## Abstract

Many fungal pathogens develop specialized infection structures such as appressoria to penetrate plant cells. However, it is not clear whether special structures are formed after cell wall penetration before invading host plasma membrane in hemibiotrophic pathogens. Here, we showed that a penetration ring consisting of Ppe1 secreted proteins is formed after appressorium-mediated cell wall penetration and remained at the base of penetration site after invading plant plasma membrane by the rice blast fungus *Magnaporthe oryzae*. The same persistent Ppe1 ring is formed after the penetration of neighboring cells by transpressoria. *PPE1* is specifically expressed during plant infection and the Δ*ppe1* mutant is defective in penetration and invasive growth. Blockage of penetration peg formation impedes the development of the Ppe1 ring. Close examinations showed that the penetration ring is formed at the collar of penetration pegs between the plant cell wall and plasma membrane and it is persistent as a fixed ring even after invasive hyphae invaded neighboring cells. Furthermore, Ppe1 is a member of an expanded family of secreted proteins that are unique to fungal pathogens using extreme appressorium turgor for plant penetration. Other members of the Ppe1 family also localize to the penetration ring for anchoring on plasma membrane during plant infection. Taken together, a penetration ring consisting of a family of secreted proteins is formed between plant cell wall and plasma membrane, which may function as a novel physical structure at the interface between the tip of penetration pegs and plant plasma membrane before the differentiation of invasive hyphae.

**Significance Statement:** Like many other plant pathogens, the rice blast fungus forms melanized appressoria (specialized infection structures) to penetrate plant cells. In this study, we showed that a penetration ring is formed by penetration pegs after appressorium-mediated penetration of plant cell wall. This ring of secreted proteins is persistent at the point of penetration pegs invading plant plasma membrane to form invasive hyphae. Therefore, after the penetration of plant cell wall, the rice blast fungus forms a penetration ring that consists of a family of secreted proteins unique to pathogens using extreme appressorium turgor for penetration and may function as a physical structure for anchoring onto plant plasma membrane and developing invasive hyphae in penetrated cells.

## Introduction

*Magnaporthe oryzae*, a filamentous ascomycete, is the causal agent of the devastating rice blast disease, which poses a significant threat to global rice production (1, 2). In addition to rice, *M. oryzae* is capable of infecting a wide range of grass species, including major cereal crops such as wheat, barley, oat, and millet, thus compromising global food security (3, 4). As one of the most important plant diseases, it is mainly controlled by planting resistant cultivars. Nevertheless, outbreaks of rice blast still occur throughout rice growing regions worldwide because of breakdown of resistance by new races of the pathogen. In addition, wheat blast caused by *M. oryzae* pathovar *Triticum* has recently spread from South America to Asia, threatening wheat production (5, 6).

Like many other foliar fungal pathogens, penetration of the cuticle and cell wall of host plants is the most critical infection step (7, 8). The infection cycle of *M. oryzae* starts with attachment of dispersing conidia to plant surface by releasing sport tip mucilage. Germ tubes emerging from attached conidia can sense surface hydrophobicity and hardness and develop highly specialized infection structures known as appressoria at the tip (1, 9). Mature dome-shaped appressoria are anchored on rice surface with appressorium mucilage (10) and heavily melanized by depositing a melanin layer between the cell wall and membrane as an impermeable barrier for turgor generation (11). Subsequently, the appressorium uses the extremely high turgor pressure to physically penetrate the cuticle and cell wall with a narrow penetration peg (7, 12, 13). As an hemibiotrophic pathogen, after the penetration of plant cell wall, the penetration peg does not penetrate plant plasma membrane but differentiates into primary invasive hyphae and invasive hyphae that are enveloped by plant-derived extra-invasive hyphal membrane (EIHM) in penetrated cells (7, 14). To date, it is not clear whether there is any special structure at the contact point between the penetration peg and plant plasma membrane before the differentiation of primary invasive hyphae. However, a biotrophic interfacial complex (BIC), a plant membrane-rich structure, is formed at the tip of the primary invasive hyphae for effector delivery (15, 16). Following proliferation within the first colonized cell, the tips of invasive hyphae differentiate into transpressoria, which subsequently form narrow infection pegs to traverse the plasmodesmata (PD)-rich cell junctions, allowing entry into adjacent host cells (17, 18). The fungus continues this colonization strategy until disease symptoms manifest on infected rice plants.

To suppress plant defense responses and facilitate invasive growth, *M. oryzae* deploys effectors with various functions that can be delivered into host tissues or cells with two secretion systems during infection (19). Apoplastic effectors, such as Bas4 and Slp1, accumulate in the extracellular space between the fungal invasive hyphal cell wall and EIHM, outlining the entire invasive hyphae, are secreted through the conventional endoplasmic reticulum (ER)-Golgi-dependent pathway which is sensitive to brefeldin A (BFA) (19). Conversely, cytoplasmic effectors, including Pwl2, AvrPiz-t, and Bas1, accumulate in the BIC before being delivered into plant cells using a Golgi-independent pathway involving the exocyst complex and t-SNAREs, which is insensitive to BFA (19). To date, all the known proteinaceous effectors in *M. oryzae* belong to these two categories and localize to the extracellular space outlining invasive hyphae or the BICs and intracellular targets inside plant cells.

In a previous study, we identified a family of small secreted proteins that may function as effectors in *M. oryzae* (20). In this study, we found that one of them named as *PPE1* (for <u>p</u>eg <u>p</u>eriphery-localized <u>e</u>ffector <u>1</u>, MGG_16603) is specifically expressed during plant infection and is a crucial virulence factor. Surprisingly, Ppe1 preferentially localizes to the penetration site in a ring conformation that is described as the penetration ring below. The penetration ring consisting of Ppe1 proteins is formed after appressorium– and transpressorium-mediated plant penetration and remained at the point where penetration pegs transitioning into primary invasive hyphae. Ppe1 homologs are only present in fungal pathogens using extreme appressorium turgor for plant penetration. Other members of the Ppe1 family also localize to the penetration ring that is at the base of primary invasive hyphae outside plant plasma membrane. Taken together, a penetration ring consisting of secreted Ppe family proteins is formed between plant cell wall and plasma membrane, which may function as a novel physical structure at the interface between the tip of penetration pegs and plant plasma membrane before differentiation into invasive hyphae.

## Results

### Ppe1 is important for full virulence of *M. oryzae*

We previously identified 21 host-adapted effector genes in *M. oryzae* that may play critical roles during rice infection based on *in silico* analyses (20). To explore the functions of these effectors, we selected *PPE1* (MGG_16603) for further functional characterization because its high transcription level during early infection stages as reported (20, 21). We first assayed the expression profile of *PPE1* during infection by qRT-PCR. The result showed that its expression increased during the early stages of plant infection (24-36 hpi) and peaked at 36 hpi (Fig. 1A). Moreover, fluorescence signals of mCherry driven by the *PPE1* promoter indicated that its expression was only observed in matured appressoria and during infection (Fig. 1B). These results were consistent with the transcriptome data, suggesting that Ppe1 is functional during appressorium penetration and plant infection. Ppe1 encodes a 116-aa protein with typical characteristics of fungal effectors, including an N-terminal signal peptide and eight cysteine residues (Fig. S1A). The signal peptide of Ppe1 is functional in directing the secretion of yeast Suc2 invertase in a yeast signal trap system (Fig. S1B), suggesting that Ppe1 is a secreted protein.

**Fig. 1.**
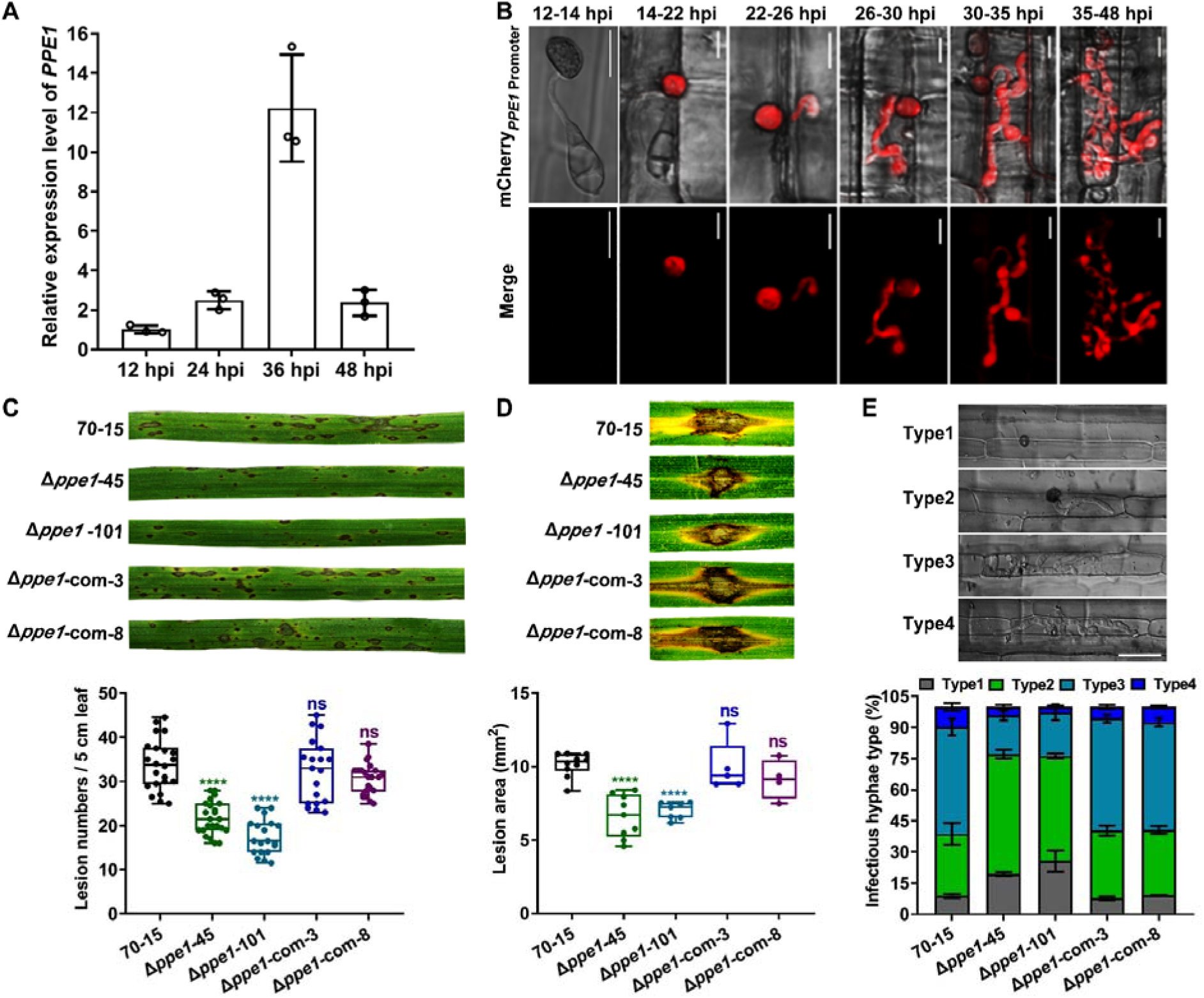
Ppe1 is important for full virulence in *M. oryzae*. (A) The relative expression level of Ppe1 during infection of rice leaf sheaths assayed by qRT-PCR. (B) Fluorescence signals of the P*_PPE1_*-mCherry transformant in penetration assays with rice leaf sheaths. Scale bars = 10 μm. (C) Typical leaves of CO-39 seedlings sprayed with conidia of the wild type (70–15), *PPE1* gene deletion mutants (Δ*ppe1-45*, Δ*ppe1-101*) and complement strains (Δ*ppe1-com-3*, Δ*ppe1-com-8*) examined at 7 days post inoculation (dpi). Boxplot below shows the number of disease lesions per 5 cm of individual inoculated leaf. n = 22, 23, 19, 19, and 22, respectively. (D) Wounded leaves of CO-39 drop-inoculated with the marked strains were examined at 10 dpi (upper panel) and measured for the lesion area (lower panel). n = 10, 9, 7, 5, and 4 respectively. (E) Rice leaf sheath penetration assays. Conidia suspensions from each strain were respectively injected into 4-5 weeks old rice leaf sheaths. Infection levels were observed at 32 hpi (hours post inoculation). The infection levels were grouped into four types: Type 1, appressorium formation without penetration; Type 2, having penetration peg or primary invasive hyphae; Type 3, having more than two branched invasive hyphae restricted in the first infected rice cell; Type 4, having invasive hyphae crossing to neighboring host cells. Scale bar = 10 μm. Statistical analyses of the infection types and percentages are also shown. Significant differences were determined by one-way ANOVA. Asterisks indicated significant differences (p<0.0001). ns indicated no significant difference (p>0.05).

To determine the biological functions of Ppe1 in *M. oryzae,* we generated a *PPE1* gene deletion mutant in the wild-type strain 70-15 (Fig. S1C). Deletion of *PPE1* had no obvious effect on the fungal growth, conidiation, and appressorium formation (Fig. S1D-G). However, in spray infection assays with rice leaves, the Δ*ppe1* mutant was significantly reduced in the number of disease lesions compared to the wild type (Fig. 1C). In wound infection assays, the Δ*ppe1* mutant also was significantly reduced in virulence and formed smaller lesions than the wild type (Fig. 1D). To confirm the role of *PPE1* in virulence, we generated a complemented strain Δ*ppe1-* com that had a copy of the *PPE1* gene ectopically integrated into its genome. In spray or wound infection assays, Δ*ppe1-*com was similar with the wild type in virulence (Fig. 1C-D), indicating a complete recovery in phenotypes. We then conducted rice leaf sheath penetration assays to observe the infection processes. At 32 hpi, over 91% of the appressoria formed by the wild type and the complemented strain had penetrated and developed invasive hyphae in rice cells.

Invasive hyphae in most of the penetrated cells (approximately 60%) had more than two branches, and approximately 10% of them had spread into the neighboring cells (Fig. 1E). In contrast, approximately 25% of the appressoria formed by the Δ*ppe1* mutant had no penetration. Although 53.5% of these mutant appressoria successfully penetrated into host cells, only 23% of Δ*ppe1* invasive hyphae had more than two branches and less than 4% had spread into adjacent cells (Fig. 1E). To further validate these observations, we generated a *PPE1* knockout in the wild-type reference strain Guy11. Similarly, we observed a significant decrease in the pathogenicity of the Δ*ppe1* mutants compared to Guy11 in the spray assay (Fig. S2). These results indicate that the Δ*ppe1* mutant has defects in appressorium penetration, invasive hyphal growth and subsequent spread of invasive hyphae into neighboring plant cells.

### Ppe1 forms a ring at the base of appressoria during host penetration

To determine the localization of Ppe1 during infection, we expressed a full-length Ppe1-mCherry fusion protein under the control of *PPE1* promoter in both the wild type (70–15) and the Δ*ppe1* mutant. Rice leaf sheaths were then infected with these strains and the localization of Ppe1 was analyzed by confocal microscopy. In these transformants, Ppe1-mCherry signals were observed as a ring underneath the appressoria (likely at the site of the emergence of penetration pegs) at 22 hpi (Fig. 2A, Video S1). The Ppe1-mCherry ring persisted after appressorial penetration and the ring-shaped fluorescence signal of the Ppe1-mCherry was still observed even after the formation of primary and secondary invasive hyphae (Fig. 2B). We further measured the size of this ring and found that the average diameter of the inner circle of the Ppe1-mCherry ring was 1.53 μm. The width of the Ppe1-mCherry ring varied from 1.79 to 4.78 μm, which is smaller than the width of appressoria (Fig. 2C) and known septin ring (5.9 μm) at the appressorium pore (13). Furthermore, the Ppe1-mCherry fusion protein could be detected by western blot analysis with proteins isolated from infected rice leave (Fig. S3). Because the Ppe1 ring always appears at the point of emergence of the penetration peg, we named this ring structure as penetration ring.

**Fig. 2.**
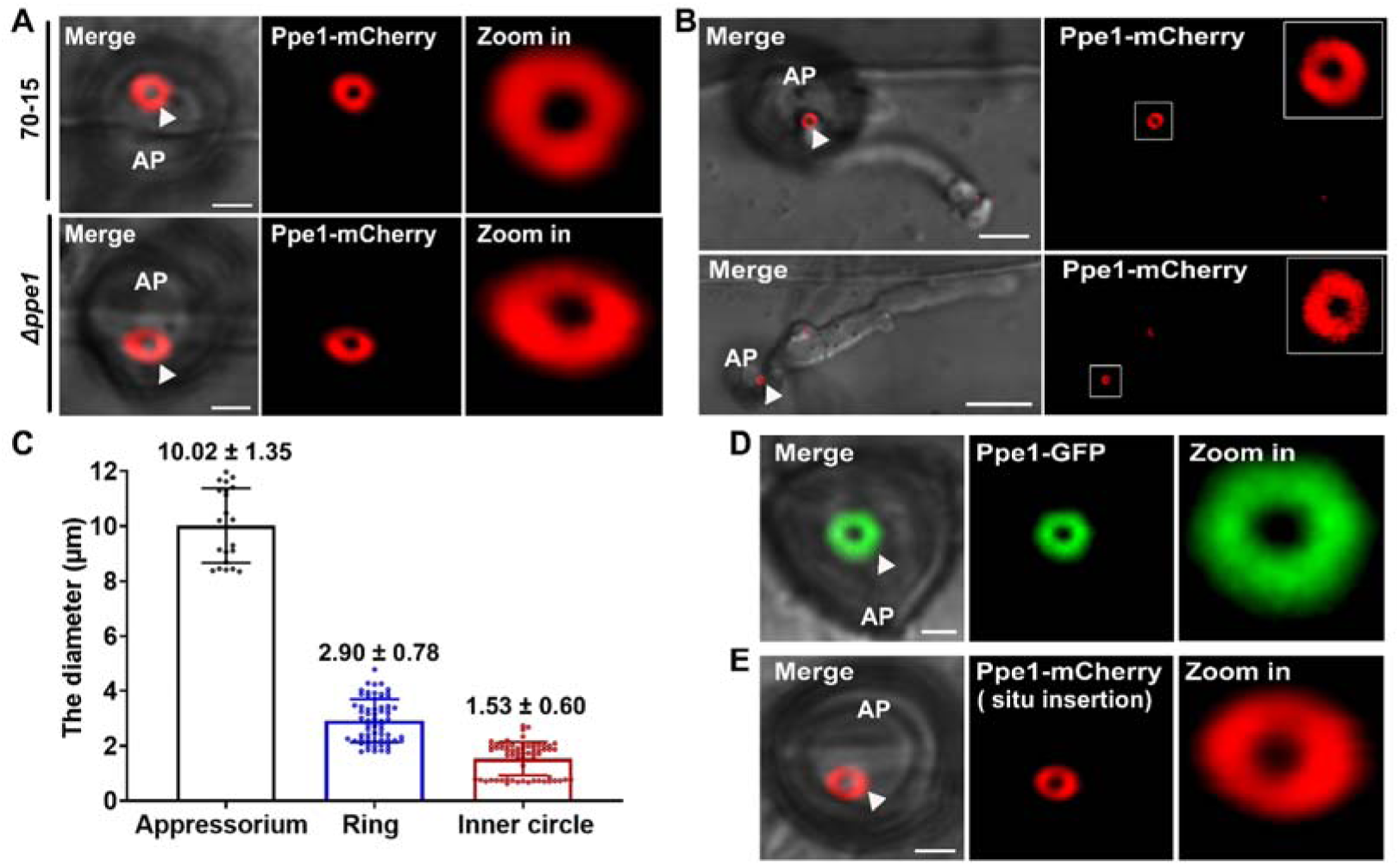
Ppe1-mCherry forms a ring during appressorial penetration. (A) A ring of Ppe1-mCherry signals at the penetration site was observed at 22 hpi in leaf sheath penetration assays with the *PPE1*-mCherry transformant. Upper panel indicates the Ppe1-mCherry expressed in wild-type strain (70–15). Lower panel indicates the Ppe1-mCherry expressed in Δ*ppe1* mutant. Scale bars = 2 μm. (B) The Ppe1-mCherry penetration ring was persist even after the growth of primary and secondary invasive hyphae. Scale bars = 5 μm. (C) The diameters of appressoria, Ppe1-mCherry penetration ring (ring), and its inner circle. n= 21, 65 and 65 respectively. (D) The formation of Ppe1-GFP rings at the penetration site formed by the *PPE1*-GFP transformant. Scale bars = 2 μm. (E) The in-situ *PPE1*-mCherry transformant also formed a fluorescent ring at the penetration site in leaf sheath penetration assays. Scale bar = 2 μm. AP, Appressorium. Triangle arrows pointed to penetration rings.

To confirm the localization of Ppe1, we also generated Ppe1-GFP transformants. Like Ppe1-mCherry, Ppe1-GFP also formed a ring at the penetration site underneath the appressorium (Fig. 2D). These results indicate that, irrespective of being fused with mCherry or GFP, Ppe1 localizes to the penetration ring. To avoid possible overexpression or unexpected effects due to ectopic integration of the fusion constructs, we generated a transformant from 70-15 strain in which mCherry was inserted right before the TAA stop codon of the endogenous *PPE1* gene. In the resulting transformant expressing the in situ *PPE1*-mCherry fusion construct, the penetration ring was also observed (Fig. 2E). These results further verify the localization of Ppe1 to the penetration ring during appressorial penetration.

### Penetration ring is also formed during the penetration of neighboring cells by transpressoria

In *M. oryzae*, appressoria-like structures called transpressoria are formed by invasive hyphae on the wall boundary of the invaded cell in order to penetrate into neighboring cells (17, 18). Therefore, we observed localization of Ppe1 during transpressorium-mediated penetration of plant cells. In the *PPE1*-mCherry transformant, like during appressorial penetration, the penetration ring was observed during transpressoria-mediated penetration of neighboring cells (Fig. 3A). Similarly, we observed the penetration ring in the neighboring cells penetrated by invasive hyphae of the *PPE1*-GFP transformant (Fig. 3B). The Ppe1-mCherry or Ppe1-GFP ring (Fig. 3A and B – zoomed view) appeared to be at the transpressorial penetration point. These results indicate that the penetration ring is also formed during invasive hyphal cell-to-cell penetration. Furthermore, similar localization of Ppe1 to the penetration ring was observed in barley epidermal cells (Fig. 3C). Taken together, we conclude that Ppe1 localizes to the penetration ring during appressorium-mediated and transpressorium-mediated plant cell penetrations in *M. oryzae*.

**Fig. 3.**
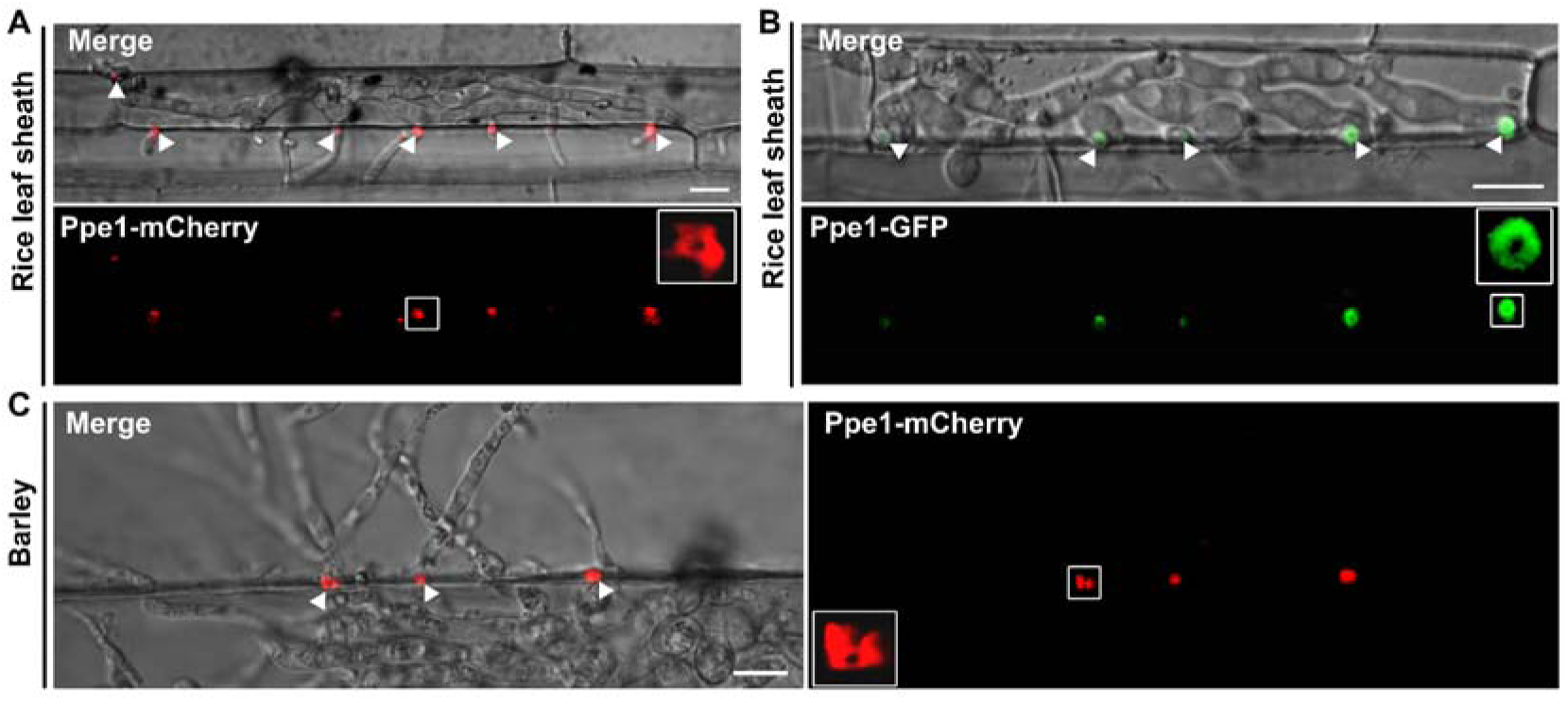
Penetration ring formation during invasive hyphal penetration into neighboring cells. (A) Ppe1-mCherry forms rings at about 36 hpi. (B) Ppe1-GFP also forms rings similar to Ppe1-mCherry during penetration to neighboring cells. (C) Ppe1 ring formation is also observed during barley infection. Scale bars = 10 μm. Arrows heads point to penetration rings at the penetration sites of invasive hyphae. White squares indicate zoomed regions.

### Penetration ring is formed on the periphery of the penetration peg

In *M. oryzae*, appressorium-mediated penetration is a septin-dependent process in which septins aggregate and generate a hetero-oligomeric 5.9 μm ring structure at the appressorial pore to re-organize the F-actin cytoskeleton network to promote the emergence of penetration pegs (13). Following the penetration peg formation, septin ring undergoes further constriction into a smaller ring (22). To clarify the precise site of the penetration ring, we expressed Ppe1-mCherry (under the control of the *PPE1* promoter) in a strain expressing Sep3-GFP (Guy11 background) and then observed their spatiotemporal dynamics during rice leaf sheath infection. At 11 hpi, Sep3-GFP big ring was formed at the appressorium pore but the Ppe1-mCherry ring was not observed (Fig. S4A, Video S2). The Ppe1-mCherry ring signal was observed at 16 hpi, after the constriction of Sep3-GFP ring and the former was completely surrounding the latter (Fig. S4B). Subsequently (at 19 hpi), the Ppe1-mCherry ring (still surrounding the Sep3-GFP ring) became constricted (Fig. 4A, Fig. S4C, Video S3). However, on impenetrable hydrophobic coverslips, the Sep3-GFP ring was formed normally at the appressorium pore, but the Ppe1-mCherry ring and constriction of Sep3-GFP were not observed (Fig. S5). These results suggest that Ppe1-mCherry ring is formed after the formation of penetration pegs, outside the constricted Sep3-GFP ring.

**Fig. 4.**
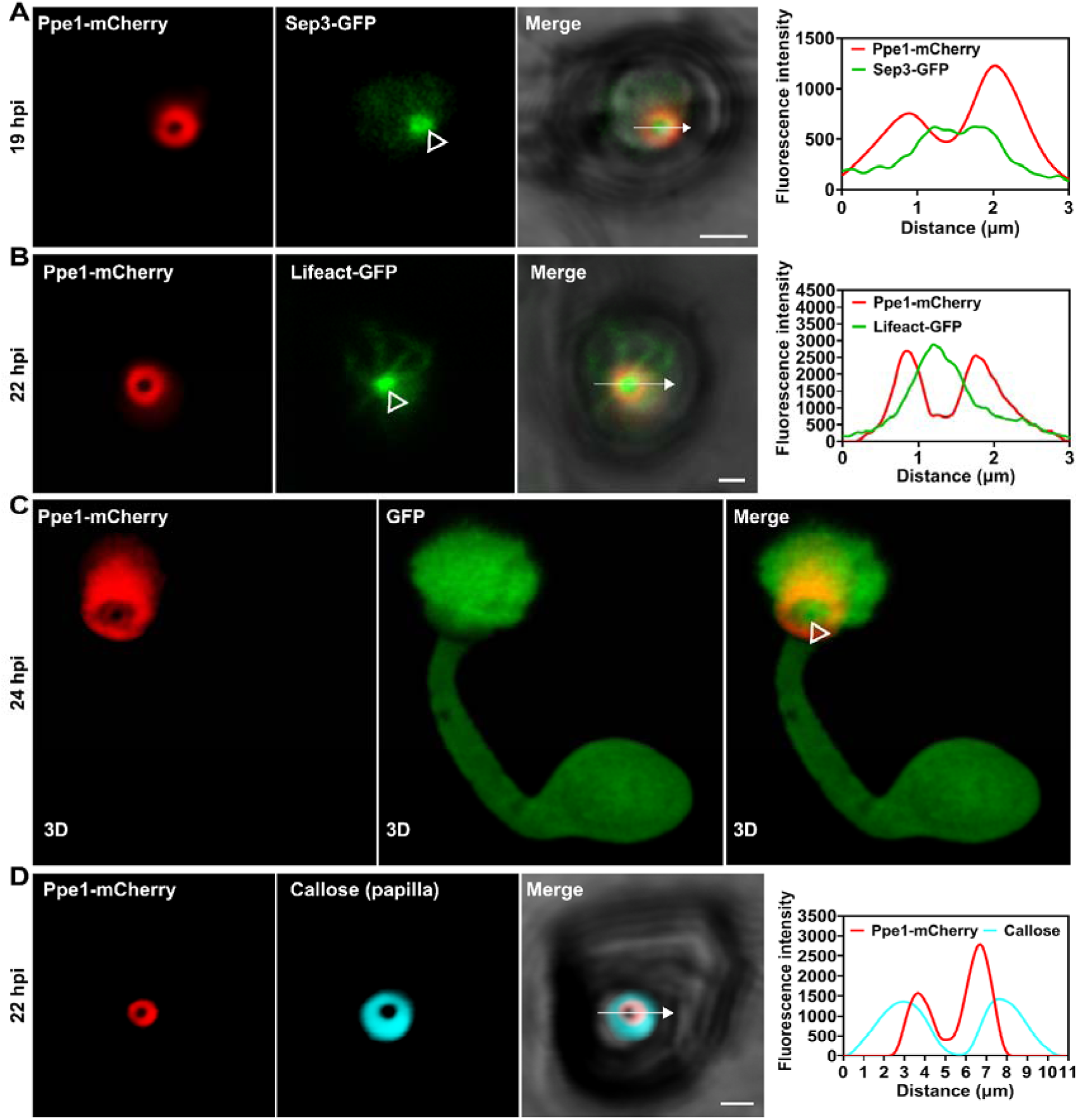
The penetration ring localizes on the periphery of penetration peg. (A) Ppe1 ring surrounds the Sep3 constricted ring. Scale bar = 2 μm. (B) Ppe1 ring also surrounds the Lifeact ring. Scale bar = 1 μm. (C) Ppe1 ring is formed around the narrow site between an appressorium and invasive hypha. Three-dimensional (3D) volume rendering images were constructed by confocal optical z-series images captured at 0.35 μm intervals, with a total of 22 steps. (D) Ppe1 ring partly colocalized with callose ring at the apoplastic papilla. Scale bar = 1 μm. Hollow triangles indicate the site of penetration peg. Line graphs were generated at the directions pointed by the white arrows.

Penetration pegs are rich in microfilaments, filasomes and microtubules (23), therefore, it can be labeled by Lifeact-GFP. Hence, we co-expressed Ppe1-mCherry with Lifeact-GFP in 70-15 and observed their positioning in penetration assays with rice leaf sheaths. The Ppe1-mCherry ring was also observed to surround the Lifeact-GFP signal (Fig. 4B, Fig. S6, Video S4), indicating that the penetration ring is localized at the periphery of penetration pegs during the initial infection stage and primary invasive hyphal development. To confirm this observation, we co-expressed the Ppe1-mCherry with cytosolic GFP in 70-15. Using a three-dimensional (3D) volume rendering image constructed by confocal optical z-series images, we found that the Ppe1-mCherry ring was situated around the narrow site between the appressorium and invasive hyphae labeled with GFP (Fig. 4C, Fig. S7).

A previous study showed that callose deposition is dynamic during *M. oryzae* infection and has three different forms: callose-rich papillae underneath appressorium penetration sites, callose collars formed at cell wall crossing sites around the invasive hyphae and callose deposition at plasmodesmata pit fields (24). To investigate whether the penetration ring locates outside the peg, we stained callose deposits with aniline blue to examine the possible positioning of the callose ring underneath the appressorial penetration site and Ppe1-mCherry. We discovered that the Ppe1-mCherry ring was surrounded by the callose ring, exhibiting partial colocalization and close attachment (Fig. 4D, Video S5). Furthermore, we examined the localization of Ppe1-mCherry in transgenic rice expressing GFP-OsPIP2 as a plasma membrane marker (25). Our data demonstrate that the penetration ring is tightly associated with the host plasma membrane (Fig. S8). Taken together, we conclude that the penetration ring locates at the periphery of the penetration peg and anchors the peg to the host plasma membrane during the initial appressorium-mediated host penetration.

### Formation of the penetration ring at the base of penetration pegs after transpressorium penetration

To determine the precise localization of the penetration ring during the transpressoria-mediated penetration, we expressed Ppe1-mCherry (under the control of the *PPE1* promoter) in a Sep5-GFP strain and then observed their spatiotemporal dynamics during rice leaf sheath infection. The results showed that Sep5-GFP first displayed a ring structure at the swollen part of the invasive hyphal tip (Fig. S9A), similar to the septin big ring at the appressorium pore. It then exhibited a collar structure at the constricted infection peg site (Fig. S9B-C), which is consistent with a previous report by Sakulkoo (26). Meanwhile, the Ppe1-mCherry ring or collar-like ring was formed outside the Sep5-GFP collar (Fig. S9B-C), indicating that the penetration ring formed during transpressoria-mediated cell-to-cell penetration is positioned outside the penetration peg. Taken together, we conclude that the penetration ring of *M. oryzae* is localized at the periphery of penetration pegs both during appressoria– and transpressoria-mediated host penetrations.

### Penetration ring formation requires the initiation of penetration peg formation

To further characterize the relationship between the penetration ring and penetration peg, we observed the expression and localization of Ppe1-mCherry in Δ*mst12,* Δ*nox2* and Δ*nox1* mutant strains that are defective in plant penetration (27, 28). In the Δ*mst12* mutant that is defective in both penetration initiation and peg formation, the Ppe1-mCherry fluorescence signal could not be detected. In the Δ*nox2* mutant that is unable to initiate penetration peg formation, the Ppe1-mCherry ring could not be formed but displayed a disorganized mass underneath the appressorium. Interestingly, in the Δ*nox1* mutant that forms normal penetration pegs but is defective in invasive hyphal elongation, the Ppe1-mCherry ring was formed during rice leaf sheath infection (Fig. 5A), indicating that penetration peg formation is a prerequisite for the emergence of the penetration ring. To confirm this observation, the localization of Ppe1-mCherry fusion proteins was observed after inducing appressorium formation on hydrophobic coverslips and cellophane membranes. Although melanized appressoria were formed, we failed to observed the Ppe1-mCherry ring on the impenetrable coverslips (Fig. 5B). However, the penetration ring is formed after the penetration of cellophane membranes (Fig. 5C). These results indicate that penetration is necessary for the formation of the penetration ring.

**Fig. 5.**
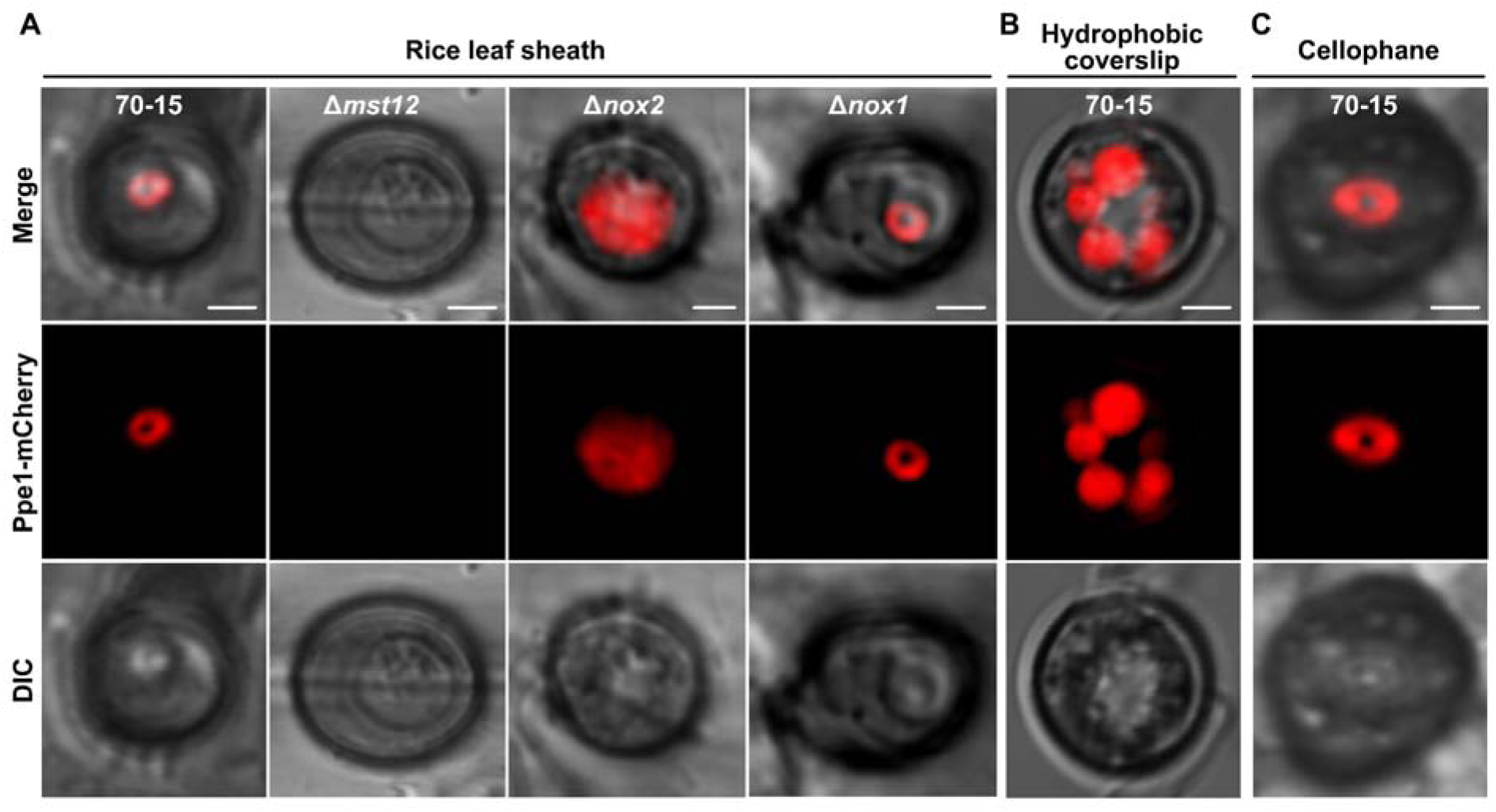
Penetration ring formation requires the emergence of penetration peg. (A) Ppe1 ring expression and localization in 70-15, Δ*mst12*, Δ*nox2* and Δ*nox1* strains at 24 hpi. Conidia suspensions from each strain were inoculated on rice leaf sheaths and observed at 24 hpi. (B) Ppe1 ring could not be formed on impenetrable coverslips. Conidia from Ppe1-mCherry strain were inoculated on hydrophobic coverslips and observed at 24 hpi. (C) Ppe1 ring was observed at 38 hpi on penetrable cellophane membrane. Scale bars = 2 μm.

### Other Ppe proteins also localize to the penetration ring

To determine whether other secreted proteins homologous to Ppe1 also localize to the penetration ring, we used the amino acid sequence of Ppe1 to carry out BLASTp search at the NCBI database to uncover the homolog proteins of Ppe1 and further analyzed their phylogenetic relationships. The Ppe1 homologs only exist in *Magnaporthe* and *Colletotrichum* species, such as the most common cereal phytopathogens *M. oryzae* and *C. graminicola* (Fig. 6A). Remarkably, these fungi are known to form melanized appressoria and deploy extremely high turgor pressure for plant penetration (8). To understand whether other members of the Ppe1 family (Ppe2, Ppe3, Ppe3, Ppe4, Ppe5, and Ppe6) from *M. oryzae* have similar localization as Ppe1, we tagged each of them with mCherry and expressed them individually in the 70-15 background under the control of their native promoters. Examination of their localizations during infection revealed that Ppe2-mCherry, Ppe3-mCherry and Ppe5-mCherry also exhibit a ring structure during appressorium-mediated host penetration, while the fluorescence signals of Ppe4-mCherry and Ppe6-mCherry were hardly observed, possibly due to their low expression (Fig. 6B). Moreover, when MoT-Ppe1 from *M. oryzae* pathovar *Triticum* was expressed in *M. oryzae* under the control of its native promoter, the characteristic penetration ring was also observed (Fig. 6B). All these results suggest that the secreted Ppe proteins accumulate at the penetration ring.

**Fig. 6.**
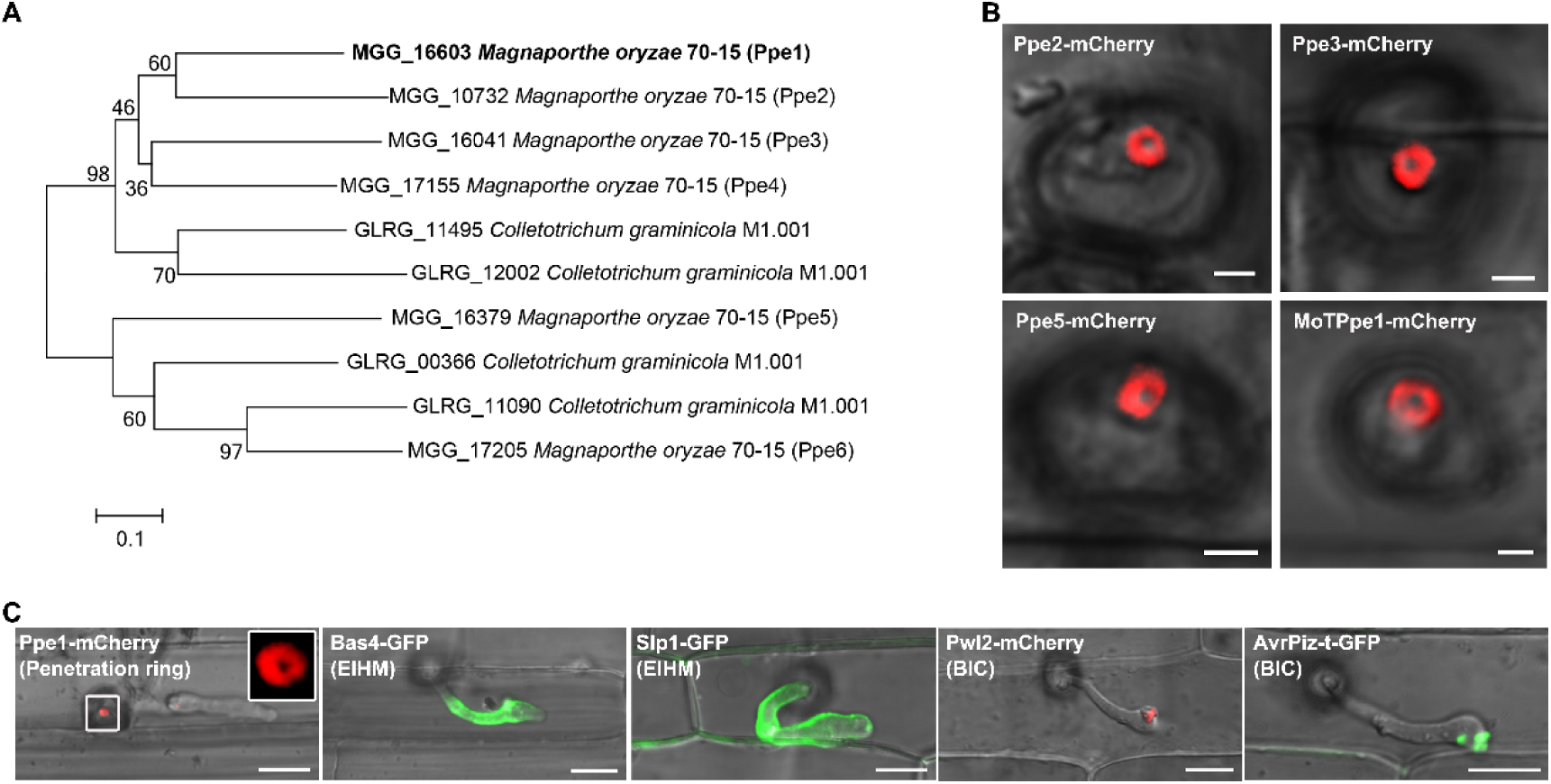
Other secreted Ppe proteins also localize to the penetration ring. (A) Phylogenetic tree of Ppe1 constructed using neighbor-joining method. (B) Paralogs of Ppe1 in *M. oryzae* localize to the penetration ring. Ppe2-mCherry, Ppe3-mCherry and Ppe5-mCherry were driven by their respective native promoters. MoTPpe1-mCherry from *Magnaporthe oryzae Triticum* was expressed in *M. oryzae* (70–15), driven by its native promoter. Scale bars = 2 μm. (C) Comparison of the localization of penetration ring with those of some previously reported fungal effectors. Bas4-GFP and Slp1-GFP uniformly outlined the extra-invasive hyphal membrane (EIHM), while Pwl2-mCherry and AvrPiz-t-GFP accumulated in the biotrophic interfacial complex (BIC). Scale bars = 10 μm.

As controls, we also examined the localizations of Bas4, Slp1, Pwl2, and AvrPiz-t. Under the same experimental conditions, Bas4-GFP and Slp1-GFP were observed to uniformly accumulate within the extra-invasive hyphal membrane (EIHM), outlining the entire invasive hyphae, while Pwl2-mCherry and AvrPiz-t-GFP accumulated in the biotrophic interfacial complex (BIC) (Fig. 6C), which is consistent with previous reports (16, 29, 30). EIHM and BIC are specifically associated with invasive hyphae during biotrophic growth. In contrast, Ppe1 localized to the penetration ring at the periphery of penetration pegs (Fig. 6C). Therefore, unlike subcellular localizations of Bas4, Slp1, Pwl2, and AvrPiz-t to the extracellular space outlining invasive hyphae, BICs, or intracellular targets in plant cells, Ppe1 and its homologs localize to the penetration ring that may function as a novel physical structure at the interface between the tip of penetration pegs and plant plasma membrane before the differentiation of invasive hyphae.

## Discussion

As the model plant pathogenic fungus, appressorium formation and appressorium-mediated penetration have been well-characterized in *M. oryzae* (18). The fungus uses extremely high turgor pressure generated in appressoria cemented to the surface with appressorium mucilage to penetrate underlying plant cells (12). As an hemibiotrophic pathogen, after piercing plant cell wall, the penetration peg does not penetrate the plasma membrane but differentiates into invasive hyphae enveloped in a layer of EIHM (16). In this study, we showed that a penetration ring of Ppe1 proteins is formed by the penetration peg on the surface of the plant plasma membrane before invading and differentiating into primary invasive hyphae (Fig. 2 and Fig. 7). Other Ppe proteins including Ppe2, Ppe3, and Ppe5 proteins also localize to the penetration ring (Fig. 6B). Because the penetration ring is persistent at the point penetration pegs transitioning into invasive hyphae outside the plasma membrane even after the penetrated cells are filled with invasive hyphae, it may function as a novel physical structure for anchoring penetration pegs on the surface of plant plasma membrane after cell wall penetration (Fig. 7). It is also possible that this penetration ring functions as a collar that is associated with the differentiation of penetration pegs (on the surface of plasma membrane) into primary invasive hyphae enveloped in the EIHM (Fig. 7). Although this transition is not well characterized, it likely involves changes in cell polarity and cytoskeleton as well as cell wall modifications. Ppe1 and its homologs may be able to assemble into a protein complex in the penetration ring, which likely involves other proteins or components as facilitating factors. In *U. maydis*, the Stp secreted effectors are known to form a protein complex during plant infection (31).

**Fig. 7.**
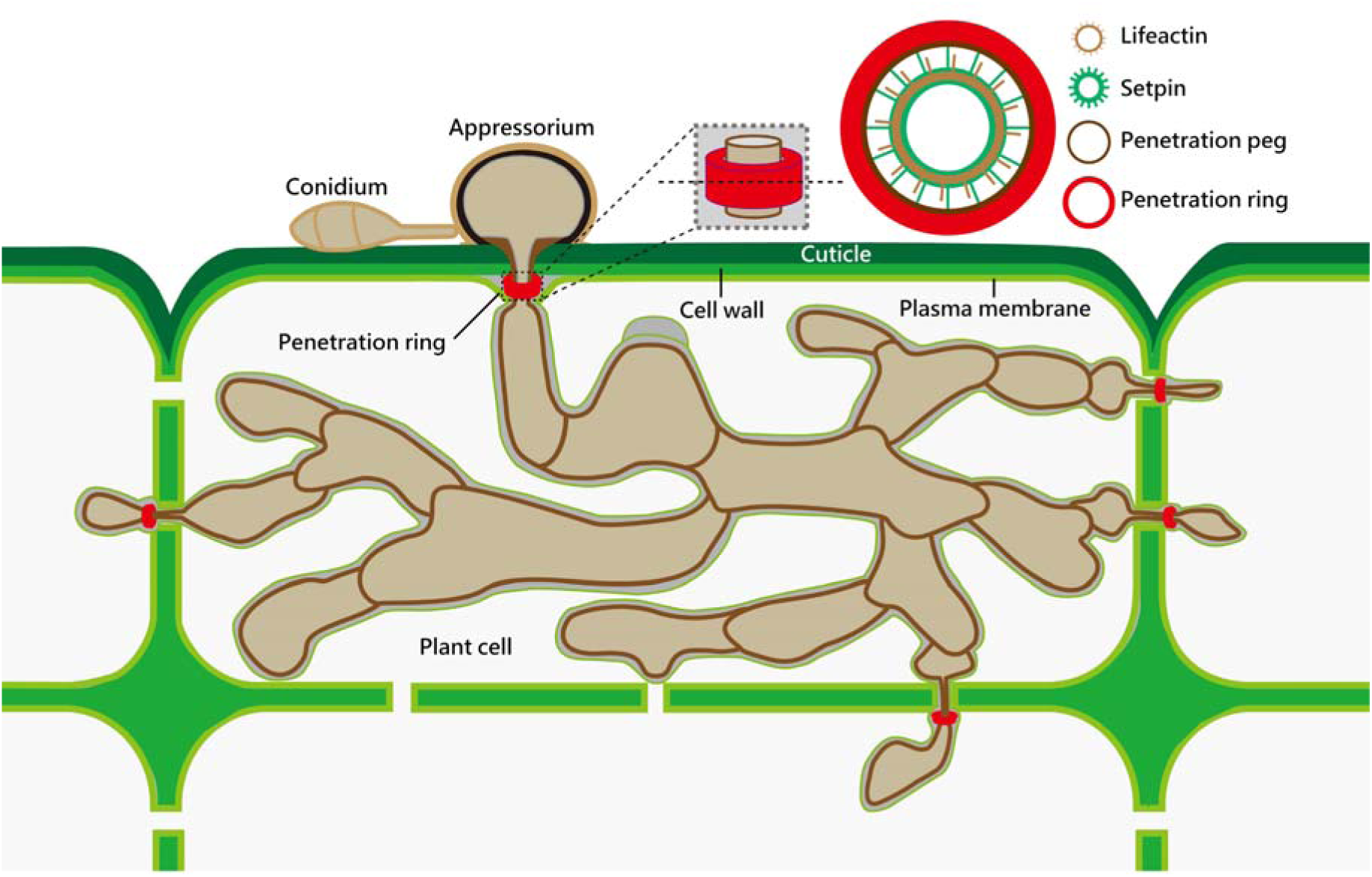
A proposed working model for the discovery of penetration ring in *M. oryzae*. Penetration ring forms at the periphery of peg and accumulates fungal secreted protein such as Ppe1 to promote *M. oryzae* invasion.

The penetration ring is also formed after transpressoria-mediated penetration of neighboring plant cells (Fig. 7). Similarly, Ppe1 and its homologs form a ring after the penetration of cell wall but before differentiating into invasive hyphae enveloped with the EIHM. The penetration ring formed by transpressoria-mediated penetration is also persistent at the penetration site on the surface of plant plasma membrane in penetrated cells (Fig. 3 and Fig. S5), suggesting that Ppe proteins are assembled into a complex that is not diffusible. In fact, similar penetration ring is observed after penetration of artificial membrane (Fig. 5C), which has no barrier to restrict the diffusion of secreted proteins, further indicating that the penetration ring of Ppe1 and other Ppe proteins is likely a structural feature or component for signaling the transition of penetration pegs to invasive hyphae. Although appressoria and transpressoria are different, the processes after the penetration of cell wall are the same in appressorium– and transpressorium-mediated plant penetration. Therefore, formation of the penetration ring in both appressorium– and transpressorium-mediated penetration is another evidence for it to be a structural element.

*M. oryzae* is known to secrete a repertoire of effectors that can be categorized into cytoplasmic and intracellular effectors (14). Ppe1 and its homologs are putative effectors that are described as HAG (host adaptation genes) effectors in an earlier study (20). Although it remains possible that Ppe1 and its homologs function as effectors during plant infection. However, in comparison with the localizations of known effectors in *M. oryzae*, we found that the localization of Ppe1 to the penetration ring is unique and Ppe1 homologs have the same localization pattern (Fig. 6). Because of the persistent localization of Ppe1 and its homologs to the penetration ring, we think the penetration ring is likely a physical structure as described above. If Ppe1 and its homologs are effector proteins being secreted at this specific location, fluorescent signals of Ppe1 and its homologs will diffuse and disappear or dimmish when effectors being translocated into plant cells. Furthermore, it is unlikely that secreted proteins localized to the penetration ring that has limited contact with the plant plasma membrane will function as effector proteins to interact with plant targets. Nevertheless, although unlikely, it remains possible that the penetration ring discovered in this study is a secretion site for secreted proteins involved in the transition of penetration pegs to invasive hyphae. After the penetration of plant cell wall, the penetration ring may not only function as a physical structure but also serve as an secretion site for the release of specific proteins to overcome plant immunity in early infection stages.

Taken together, in this study we observed the formation of the penetration ring that appears to be a structural element for signaling or triggering the differentiation of penetration pegs after the penetration of plant cell wall into invasive hyphae enveloped by the EIHM in *M. oryzae*. Based on the limited distribution of Ppe1 homologs, it is likely that only fungal pathogens using high appressorium turgor for plant penetration, such as *M. oryzae* and *Colletotrichum* species, developed the penetration ring. Therefore, it will be important to further characterize the functional relationship between the penetration ring and appressorium turgor as well as the transition of penetration pegs driven by turgor pressure into primary invasive hyphae (lower turgor pressure).

In summary, our findings have several significant implications. First, we believe that the discovery of the penetration ring as a novel physical structure associated with the differentiation of invasive hyphae represents a breakthrough in plant-pathogen interactions. Secondly, our study presents new role of the peg as a specialized platform for the deployment of secreted protein, in addition to its commonly known role as a physical penetration tool for the pathogen. Thirdly, we identified Ppe1 as a potential molecular target for controlling the devastating rice blast disease, as Ppe homologs are absent in plants and mammals. The primary challenge for future research lies in elucidating the pathogenesis associated with penetration rings. Achieving this will further our understanding of the mechanism of rice blast fungus-host interaction.

## Materials and Methods

### Strains and growth conditions

The *M. oryzae* strains 70-15 and Guy11 were used as the wild types in this study; all other fungal strains generated from the wild types are listed in Table S1 in the supplemental materials. All the fungal strains were grown on complete medium CM (CM: 10 g D-glucose, 2 g peptone, 1 g yeast extract, 1 g casamino acids, 50 mL 20 × nitrate salts (120 g/L NaNO_3_, 10.4 g/L KCl, 30 g/L KH_2_PO_4_, and 10.4 g/L MgSO_4_), 1 mL trace elements (0.1 g/L biotin, 0.1 g/L pyridoxine, 0.1 g/L thiamine, 0.1 g/L riboflavin, 0.1 g/L p-aminobenzonic acid and 0.1 g/L nicotinic acid), adjusted to pH 6.5 with NaOH, 15 g agar and distilled water added to 1 L) (32). Agar cultures were incubated at 26 °C with a 12-h light/12-h dark photoperiod for mycelial growth and conidiation. Liquid CM medium was used to harvest mycelia for extraction of genomic DNA and protoplast preparation. The bacterial strains *Escherichia coli* DH5α and *Agrobacterium tumefaciens* AGL1 were grown in Luria-Bertani (LB) medium at 37 °C and 28 °C, respectively.

### Plant and growth conditions

Rice cultivar CO-39 (*Oryza sativa* cv. CO-39) was grown in a greenhouse at 26 and 70% relative humidity with a 14-h light/10-h dark photoperiod. Barley (*Hordeum vulgare* cv. Golden Promise) was grown at 23 with a 12-h light /12-h dark photoperiod.

### Bioinformatic and phylogenetic analyses of Ppe1 homolog proteins

The Ppe1 was previously sequenced and the amino acid sequence established from our lab (33). Ppe1 signal peptide was predicted using SignalP 5.0 software. BLASTp was used to find Ppe1 homolog proteins at the NCBI database (https://www.ncbi.nlm.nih.gov/) and the representative homolog protein sequences were downloaded from the database. Phylogenetic tree was generated using MEGA7.0 software by neighbor-joining method.

### Construction of recombinant DNA and *M. oryzae* transformation

The plasmids used in the study were generated by infusion cloning method based on homologous recombination using ClonExpress II one-step cloning kit or ClonExpress MultiS kit (Cat. No. C112-01 or C113-01, Vazyme, Nanjing, China). The primers used (Table S2 in the supplemental material) were specially designed to contain 18-22-bp overlap with adjoining sequences to allow the fragments assemble by homologous recombination. To observe the expression pattern and localization of *PPE1*, the *PPE1* promoter (1485 bp) fragment amplified from the genomic DNA of the wild type (70–15) was seamlessly cloned into the *Kpn*I/*Hin*dIII-digested pKNT-mCherry/GFP fusion vectors (34) using the ClonExpress II one-step cloning kit (Cat. No. C112-01, Vazyme, Nanjing, China). For Ppe1-mCherry/GFP plasmid, the entire Ppe1 coding sequence with its native promoter but without stop codon was amplified and cloned into the *Kpn*I/*Hin*dIII-digested vector pKNT-mCherry/GFP. The pKNT-AvrPizt-GFP and other plasmids used were was constructed using the same method. The constructed plasmids were transformed into the fungal protoplasts via PEG-mediated transformation (35) and further verified by PCR screening and confocal microscopic examination of fluorescence signals. For situ fusion of Ppe1 with mCherry, the mCherry was inserted just before the Ppe1 stop codon and the construct was transformed by *Agrobacterium tumefaciens*-mediated (ATMT) transformation for homologous recombination (35).

### Assays for the secretory function of the SP of Ppe1

To confirm the secretory function of the predicted signal peptide (SP), yeast signal trap system was performed as described previously (36). The predicted SP nucleotide sequences of Ppe1 (60 bp) and Pwl2 (63 bp) were respectively cloned into pSUC2 vector that carries a truncated invertase gene *SUC2.* The generated pSUC2-SP constructs and the empty pSUC2 vector were transformed into the yeast *suc2* mutant YTK12, respectively, and then their growth conditions analyzed on CMD-W (minus Trp) plates (0.67% yeast nitrogen base without amino acids, 0.075% tryptophan dropout supplement, 2% sucrose, 0.1% glucose, and 2% agar) and YPRAA medium plate (1% yeast extract, 2% peptone, 2% raffinose, and 2 μg/mL antimicyn A) containing raffinose and lacking glucose. Transformants of YTK12 carrying the empty pSUC vector or pSUC2-Pwl2SP were used as negative and positive controls, respectively. Reduction of 2,3,5-triphenyltetrazolium chloride (TTC) to insoluble red-colored 1,3,5-triphenylformazan (TPF) was used as an indicator of the invertase enzymatic activity (37). Each of the above YTK12 transformants strain was inoculated in 5 mL of CMD-W liquid medium and incubated for 24 h at 30. The cells were then collected, washed and resuspended in 0.1% TTC in new centrifuge tubes and further incubated at 35 for 35 min. The color change was observed after 10 min of incubation at room temperature to detect the signal peptide secretion function.

### Targeted gene deletion and complementation

Deletions of *NOX1*, *NOX2* and *MST12* genes were performed using Split-marker recombination method (38). The open reading frames (ORFs) of the various genes were replaced with a hygromycin phosphotransferase (*HPH*) resistance cassette, respectively. Briefly, about 1-kb upstream and downstream sequences of the targeted genes were amplified from the genomic DNA of the wild type (70–15) using specially designed primer pairs AF+AR and BF+BR, respectively. At the same time, the upstream and downstream sequences of the *HPH* with overlapping segments were amplified from pCB1003 plasmid using the primer pairs HYG/F + HY/R and YG/F + HYG/R, respectively. The corresponding upstream and downstream fragments were respectively ligated by splicing for overlap extension (SOE) PCR using the primer pairs AF+HY/R and YG/F+BR. The PCR products were purified and transformed into the Ppe1-mCherry strain protoplasts via a PEG-mediated transformation (35). Putative transformants were selected in the presence of hygromycin B and then confirmed by PCR using the primers OF+OR and UA+H853. PCR confirmations were repeated three times for consistency. All the primer sequences used are listed in Table S2 of the supplemental material.

For *PPE1* gene deletion, *HPH* sequence was inserted between the upstream and downstream sequences of the *PPE1* gene, replacing the ORF of the *PPE1* gene, in an ATMT1 plasmid using a ClonExpress MultiS One Step Cloning Kit (Cat. no. C113-01, Vazyme, Nanjing, China). The constructed plasmid was introduced into *Agrobacterium tumefaciens* AGL1 using freeze-thaw method (39). *Agrobacterium tumefaciens*-mediated transformation (ATMT) of the construct into the wild-type spores was then carried out. The transformants were selected on hygromycin B plates and confirmed by PCR. The mutants were further confirmed by Southern blot assay using DIG-High Prime DNA labeling and detection starter kit I (Cat. no. 11745832910, Roche, Mannheim, Germany). For complementation, the constructed plasmid pKNT-Ppe1-mCherry was transformed into the Δ*ppe1* mutant protoplast and further verified by PCR screening and confocal microscopic examination for fluorescence signal detection.

### Live cell imaging

To monitor the expression and subcellular localization of Ppe1 during plant infection, rice leaf sheath inoculation and live cell imaging of fluorescently labeled proteins were performed as described (15) with slight modifications. In brief, 5-7 cm long leaf sheath segments were cut from 4-week-old CO-39 rice seedlings, and the whole hollow spaces of the sheaths were filled with conidia suspensions (5×10^4^ conidia/mL) using a 200 μL pipette. The inoculated sheaths were placed horizontally (fixed with wet filter paper fragments) on an inverted 96-well PCR plate covered by one layer of wet filter paper and incubated in a sealed box whose inner bottom was lined with two layers of wet filter paper to keep the environment humid at 26. The epidermal parts of the inoculated sheaths were trimmed into thin and transparent samples and immediately imaged using an inverted Nikon A1 laser scanning confocal microscope (Nikon, Tokyo, Japan) at the indicated time points post inoculation. The 60×/1.42 numerical aperture (NA) oil immersion objective lens was used for the imaging. The fluorescence and differential interference contrast (DIC) images were collected by confocal optical z-series acquired at 0.28 µm intervals spanning the entire fluorescence signal depth between the top plane of the appressorium formed and the bottom plane of the rice leaf sheath cross sections. The fluorescence excitation/emission wavelengths were 488 nm/500-550 nm for GFP, 561 nm/570-620 nm for mCherry and 405 nm/425-475 nm for callose staining. The images were processed using NIS-Elements AR analysis software (Nikon, Tokyo, Japan). Unless otherwise stated in the figure legend, micrograph fluorescence represents maximum intensity projection of z-series (with a 0.28 µm intervals) after removing extraneous z-planes that do not contain relevant information for a particular fluorescence. Three-dimensional (3D) volume rendering images and videos were generated from z-series in maximum intensity projection.

To observe the subcellular localizations of Ppe1 in conidia incubated on artificial hydrophobic surfaces and barley leaves, conidia suspensions (5×10^4^ conidia/mL) were dropped on the surfaces of hydrophobic coverslips or on the back sides of barley leaves (5-day old plants) and incubated under humid and dark conditions at a 26. For observation, the hydrophobic coverslips containing the appressoria were inverted, placed on 24mm×50 mm microscope glass slides and then observed under a scanning confocal microscope. Epidermal parts were removed from the infected barley leaves for observation. For penetration assay, cellophane was cut into round sheets of 4 cm in diameter and sterilized by autoclaving at 115 for 15 min. The sterile cellophane sheets were placed flat on CM plates and then the conidia suspensions (5×10^4^ conidia/mL) were dropped on the surface of the cellophane and incubated at 26. Small pieces of the cellophane containing mycelia were cut for observation under a confocal microscope at 36-48 hpi.

### Callose staining

For visualization of deposited callose, aniline blue (Cat. no. 28631-66-5, Sangon, Shanghai, China) staining was adopted (26). Rice leaf sheaths were inoculated with conidia suspensions (3×10^4^ conidia/mL). The inoculated sheaths were trimmed to thin and transparent samples at the indicated time points and stained with aniline blue (0.1mg/mL) solution prepared in 150 mM KH_2_PO_4_ (pH 9.5) for up to 30 min. The stained samples were imaged by using a laser scanning confocal microscope (Nikon, Japan).

### Phenotypic analysis

For vegetative growth and conidiation assays, small mycelial blocks from each strain were cut from the edges of 4-day-old cultures and placed at the center of freshly prepared CM plates (90 mm) and incubated for ten days at 26. The colony diameters of the cultures were measured using vernier calipers and then photographs of the cultures were taken. The cultures were then washed with 5 mL sterilized double-distilled water, filtered through three layers of lens paper and the conidia counted using a hemocytometer under a light microscope. For conidia germination and appressoria formation, spore suspensions (1×10^5^ conidia/mL) were incubated on hydrophobic coverslips (Cat. no. 12-545-101P, Fisherbrand™ Cover Glasses, USA) under humid and dark conditions at 26 for 24 h. The conidia were then observed under a light microscope for germination and formation of appressoria.

To investigate the pathogenicity of each strain, conidia spray, punch and leaf sheath inoculations were performed. Conidia suspensions (5×10^4^ conidia/mL containing 0.02% Tween-20) were used to spray the leaf surfaces of 3-week-old CO-39 rice seedlings. The inoculated seedlings were incubated in a growth chamber at 25 with 90% humidity. The seedlings were first kept in the dark for 24 h, and then subjected to 12 h light / 12 h dark photoperiod for 7 days, after which photographs were taken and the number of disease lesions on the leaf surfaces counted. Punch inoculation assay was carried out according to a method previously described (30). Briefly, 5-week-old CO-39 rice leaves were slightly wounded using mouse ear punch and 10 μL of conidia suspensions (5×10^5^ conidia/mL containing 0.02% Tween-20) from the various strains were respectively dropped at the wounded regions. Then, both sides of each inoculated spot were sealed with scotch tape to sustain the spore suspension. The inoculated plants were kept in a growth chamber under the same conditions described above for 10 days. The disease lesions formed were scanned, and later analyzed using Adobe Photoshop software to measure the lesion areas. For leaf sheaths inoculation, conidia suspensions (1×10^5^ conidia/mL) from the various strains were respectively injected into 4-5 weeks old CO-39 rice leaf sheaths. The epidermal parts of the infected leaves were trimmed to thin and transparent samples for observation under a light microscope at 32 hpi. At least 100 infected sites were observed using a Nikon A1 laser confocal microscope (Nikon, Tokyo, Japan).

### qRT-PCR analysis

To analyze the expression level of *PPE1* during *M. oryzae* infection, rice leaf sheaths were cut from 4-to 5-week-old CO-39 rice seedings and inoculated with conidia suspensions (2×10^5^ conidia/mL) from the wild type (70–15). The epidermal parts of the inoculated sheaths were cut and collected at 12-, 36– and 48 hours post inoculation (hpi). At least 15 segments of the inoculated rice sheath samples were collected at each time point. Total RNAs were extracted from the leaf sheaths or rice seeding leaves using an Eastep Super Total RNA Extraction Kit (Cat. no. LS1040, Promega, Shanghai, China). Reverse transcription was used to generate cDNA using a reverse transcription kit (Cat. no. RR037B, Takara, Dalian, China). Quantitative PCR was finally conducted using a SYBR kit (Cat. no. RR067A, Takara, Dalian, China) using specific primers (Table S2). Fugal β*-TUBULIN* gene used as endogenous reference genes, and the expression levels were calculated using 2^−ΔΔCt^ method as previously reported (40). All qRT-PCR assays were repeated three times for each sample.

## Statistical analyses

Statistical analyses were performed using GraphPad Prism7 (GraphPad Software, Inc., San Diego, CA). Statistically significant differences were determined by one-way ANOVA with two-sided least significant difference. **** indicated significant difference at p < 0.0001, ns indicated no significant difference.

## Data Availability

The authors declare that all the data supporting the findings of this study are available within the paper and SI Appendix. Raw data and details of experimental procedures are available upon request from the corresponding authors.

## Acknowledgments

We thank Prof. Nicholas J. Talbot and Prof. Barbara Valent for providing us with Δ*sec5* mutant and pBV591 plasmid respectively. We also thank Prof. Min Guo for providing us with Sep3/5-GFP strains and Δsep3/4/5 mutants. We also appreciate Profs. Guodong Lu, Jie Zhou, Huawei Zheng, Wei Tang, Ya Li, Xiaofeng Chen, and Huakun Zheng, as well as Drs Yijuan Hang, Meilian Chen and Yonghe Hong for their helpful suggestions. Thanks to Jin Zhang, Chunling Li and Yuping Fan’s help. This research was supported by the National Key Research and Development Program of China (2023YFD1400200), the National Natural Science Foundation of China (32122071, 31601596), and the Natural Science Foundation of Fujian Province (2021J06015).

## Supplementary figures

**Fig. S1.**
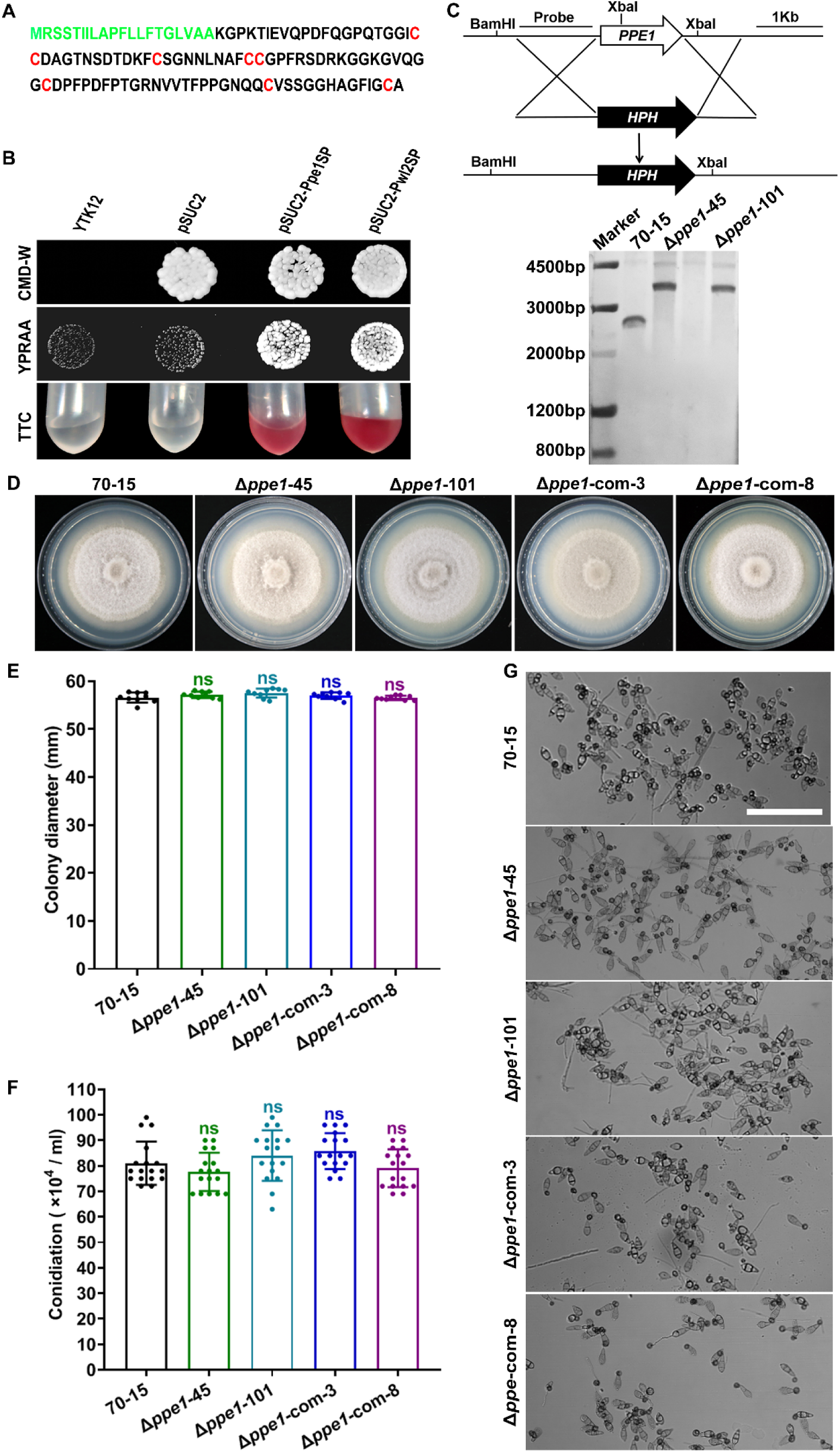
Ppe1 is dispensable for mycelial growth, conidiation and appressorium formation in *M. oryzae*. (A) Amino acid sequence of *M. oryzae* Ppe1. Signal peptides (green) were predicted by Signal-5. Cysteine residues are shown in red. (B) Detection of Ppe1 signal peptide secretion by yeast signal trap assay. The recombinant pSUC2-Ppe1SP and pSUC2-Pwl2SP constructs were transformed into the yeast strain YTK12 respectively. The pSUC2-Pwl2SP-expressing strain served as a positive control while pSUC2 empty vector was used as a negative control. (C) Targeted gene deletion strategy for *PPE1* in *M. oryzae* 70-15. Schematic map showing targeted disruption of *PPE1* gene via Agrobacterium tumefaciens-mediated transformation. The Southern blot result shows a successful replacement of *PPE1* by single insertion of hygromycin phosphotransferase (*HPH*) resistance cassette at *PPE1* ORF locus. (D) Colonies of the wild type (70–15), two independent Δ*ppe1* transformants, and complemented strains. (E) Statistical analyses of the colony diameters of each strain after being cultured on complete medium (CM) at 25 °C for 10 days. n = 9. (F) Statistical analyses of the number of conidia produced by each strain after being cultured on CM at 25 °C for 10 days. n = 17. (G) Appressorium formation by the indicated strain observed at 24 hours on artificial hydrophobic surfaces. Scale bar = 50 μm. Significant differences were determined by one-way ANOVA. ns stands for no significant difference (p>0.05).

**Fig. S2.**
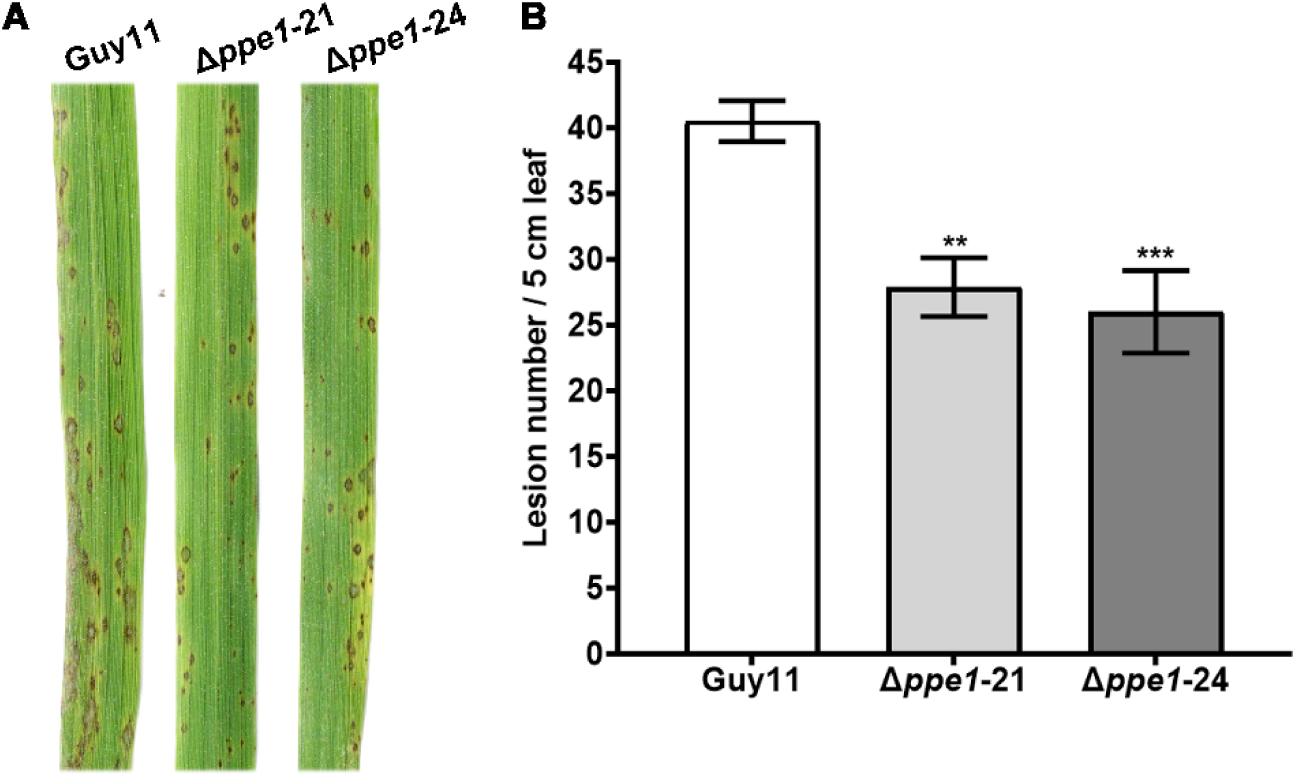
Ppe1 is essential for virulence in *M. oryzae* strain Guy11. (A) Disease symptoms on leaves of CO-39 rice seeding inoculated with conidial suspension (5× 10^4^ spores/ml) from Guy11, Δ*ppe1-21* and Δ*ppe1-24*. Photographs were taken 7 days after inoculation. (B) Quantification of the lesion number per 5 cm length of rice leaf. Asterisk represents significant difference. **, p < 0.01. ***, p < 0.001.

**Fig. S3.**
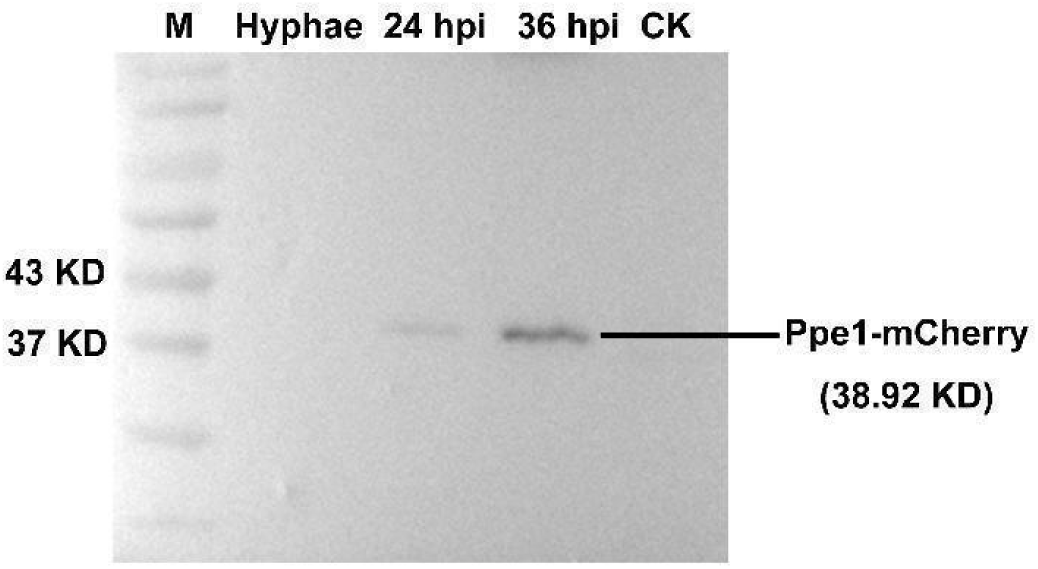
Detection of Ppe1-mCherry fusion protein by western blot. M, protein marker. CK, rice leaf sheaths injected with sterile double distilled water.

**Fig. S4.**
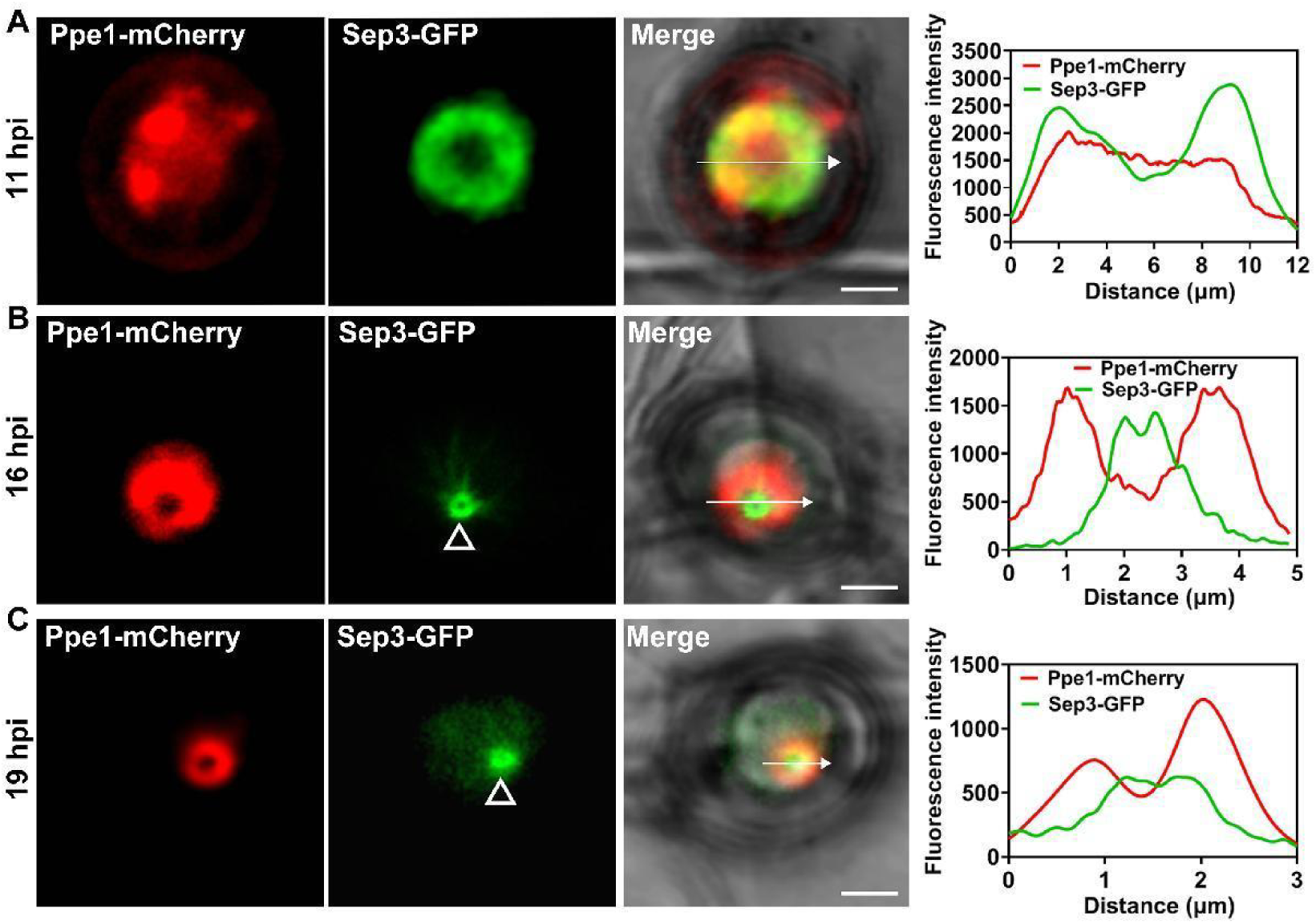
Ppe1 ring is formed simultaneously with peg penetration. (A-C) Spatiotemporal dynamics of Ppe1 ring and Sep3 ring during appressorium-mediated peg penetration into rice leaf sheaths. Scale bars = 2 μm. Hollow triangles show the sites of penetration peg. Line graphs were generated at the directions pointed by the white arrows.

**Fig. S5.**
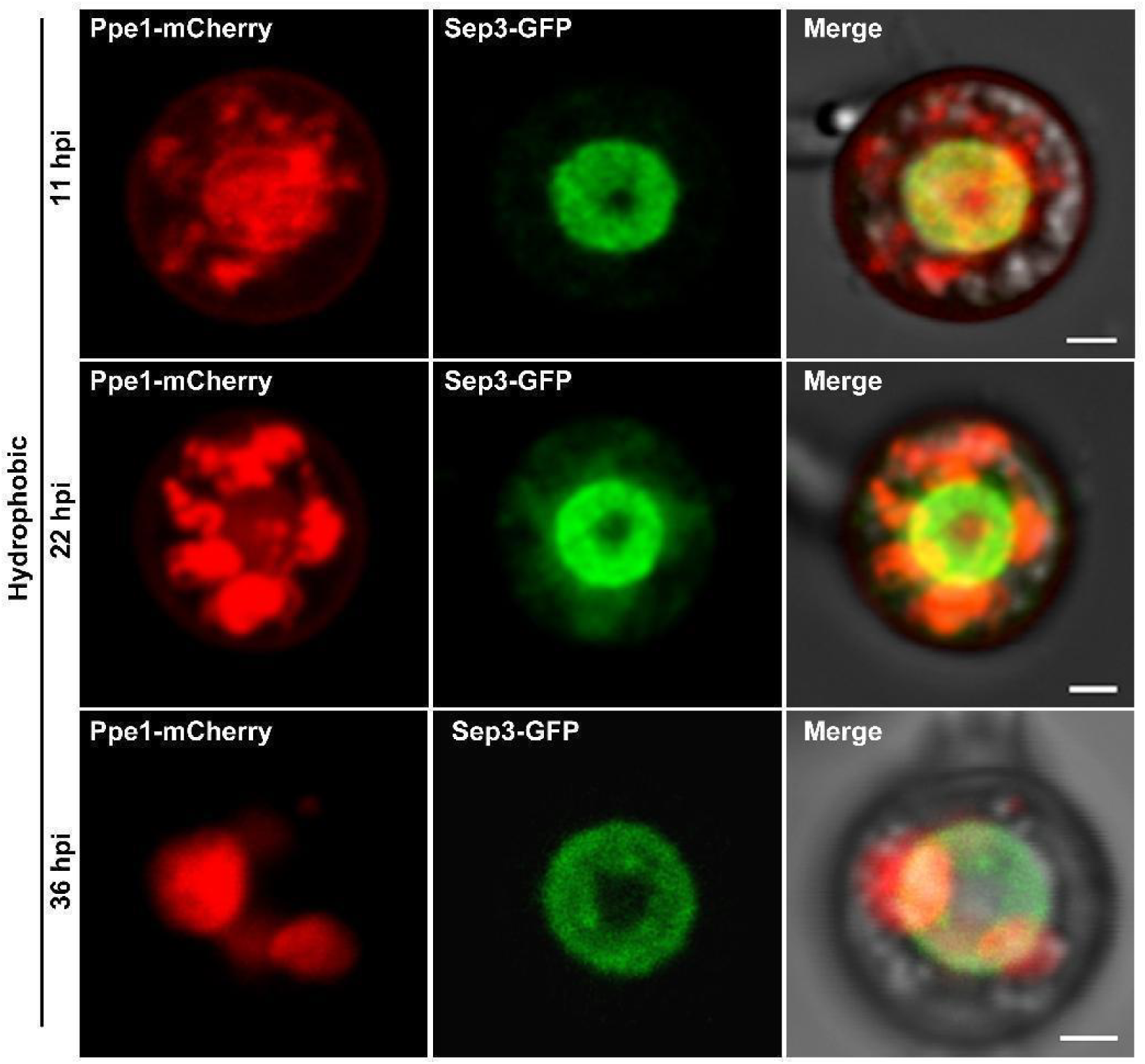
Spatiotemporal dynamics of Ppe1-mCherry and Sep3-GFP on hydrophobic coverslips. Conidia were harvested from a strain co-expressing Ppe1-mCherry and Sep3-GFP and inoculated on hydrophobic coverslips and observed at each time point. Scale bars = 1 μm.

**Fig. S6.**
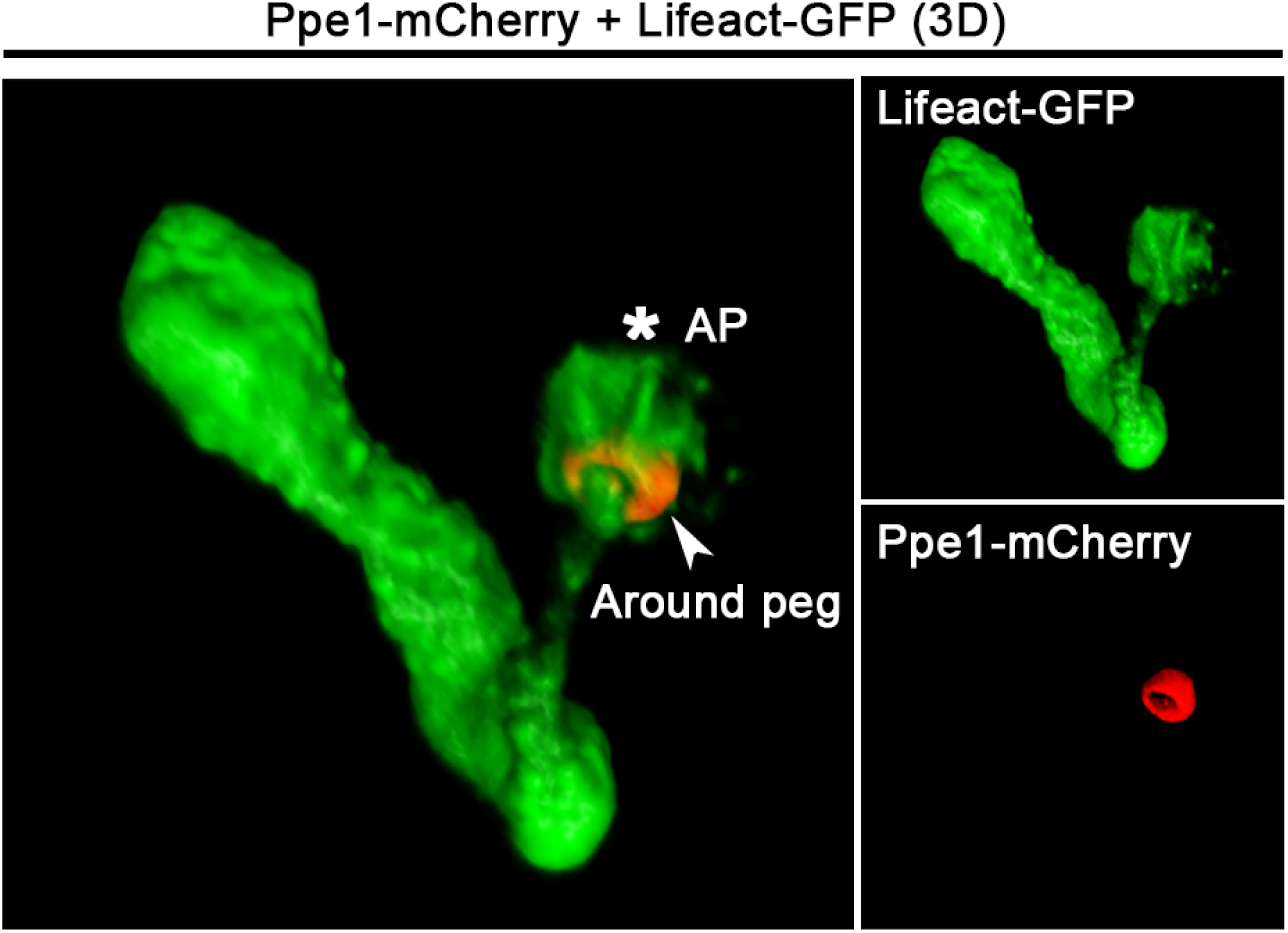
3D images of the co-expression of Ppe1-mCherry and Lifeact-GFP during primary invasive hyphal development. AP, Appressorium. Arrow shows penetration ring.

**Fig. S7.**
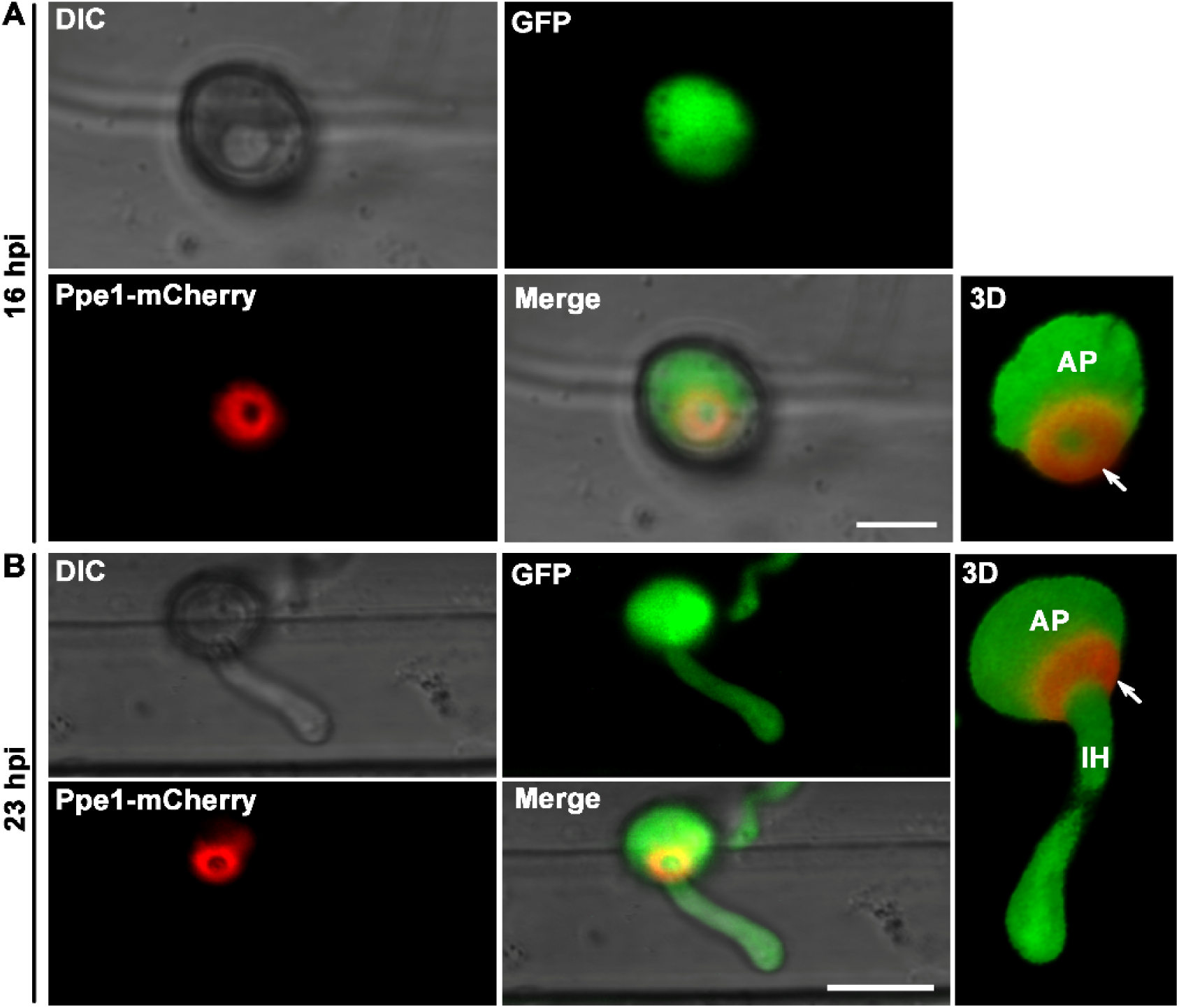
Co-expression of Ppe1-mCherry with cytoplasmic GFP at the early infection stage of *M. oryzae*. (A) Ppe1-mCherry signal formed a tight ring-like structure closely associated with the base of the appressorium at 16 hpi. Scale bar = 5 µm. (B) At 23 hpi, Ppe1-mCherry ring signal was still beneath the appressorium, and around the penetration peg which differentiated into the primary invasive hyphae. Scale bar = 10 µm. AP, Appressorium. IH, invasive hyphae. hpi, hours post-inoculation. Arrows show penetration ring.

**Fig. S8.**
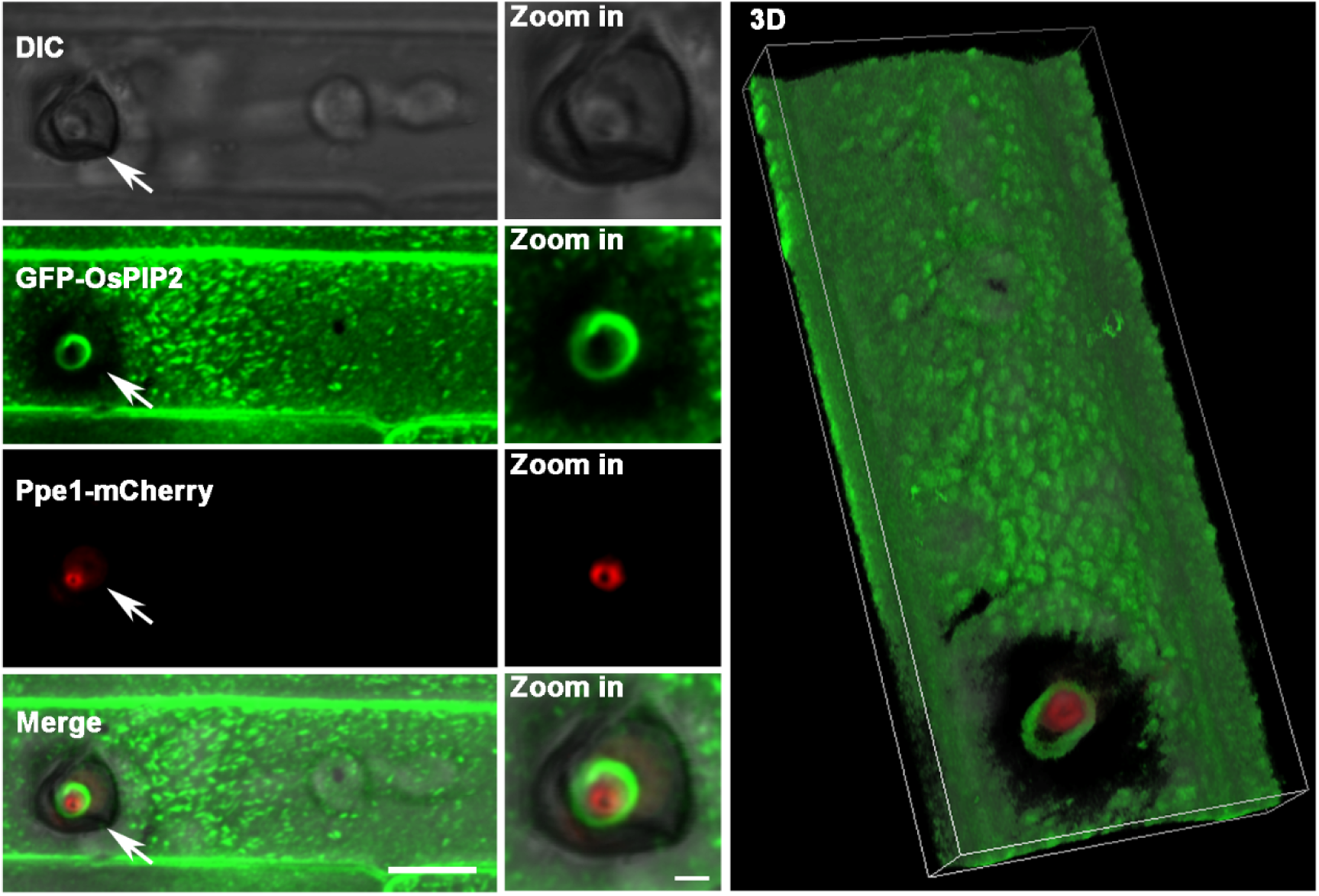
The spatial positioning of Ppe1-mCherry ring and rice plasma membrane (marked by GFP-OsPIP2). The zoomed images represent the sites pointed by the white arrows, respectively. Scale bar = 10 µm; 2 µm for the zoomed images.

**Fig. S9.**
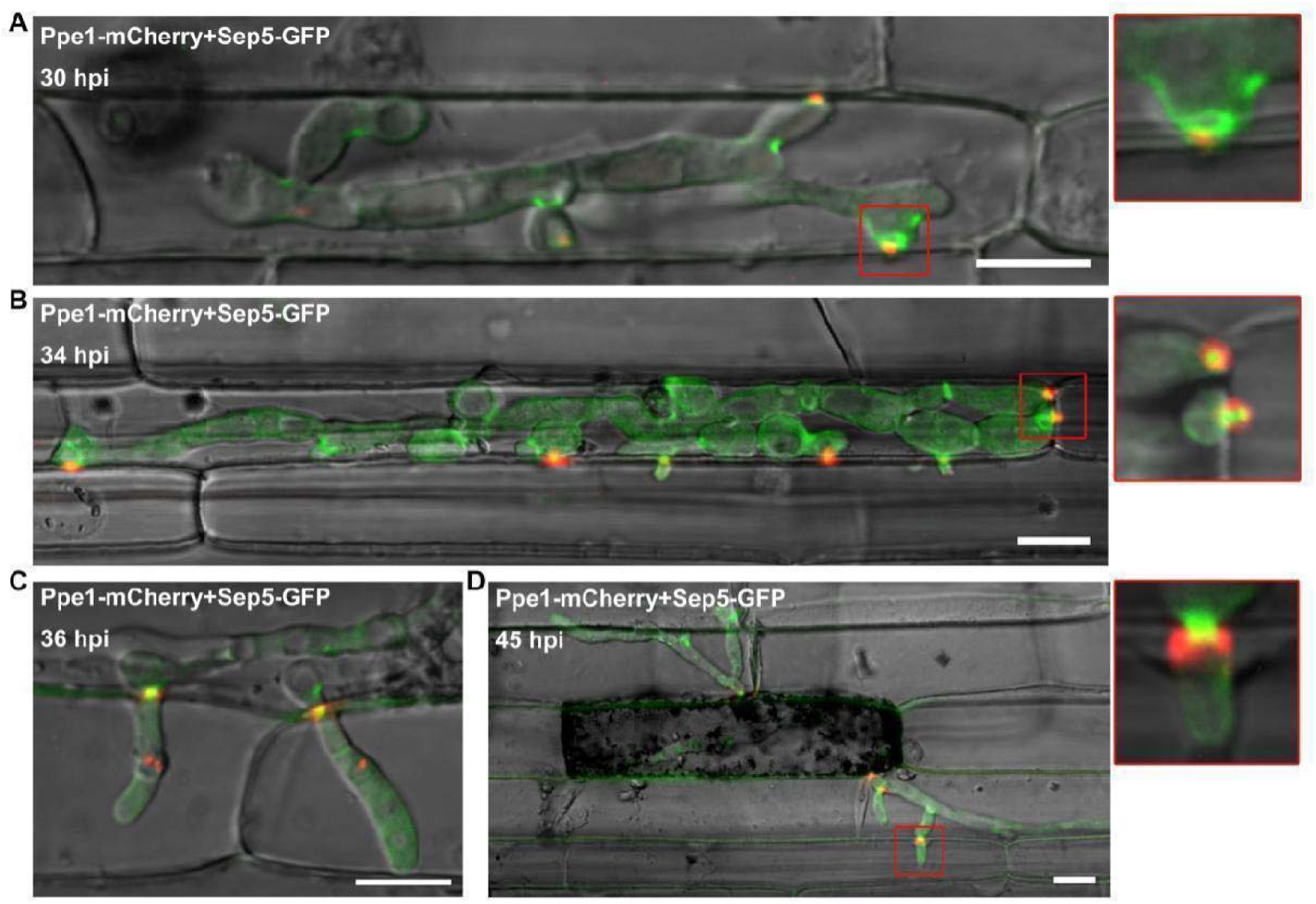
Ppe1 ring was also observed during transpressorium-mediated peg penetration. (A-D) The spatiotemporal dynamics of Ppe1-mCherry and Sep5-GFP during transpressorium-mediated cell-to-cell penetration. Red squares indicate the zoomed regions, respectively. Scale bars = 10 μm.

## Supplementary Videos

Video S1. 3D visualization of Ppe1-mCherry ring formation at 22 hpi.

Video S2. 3D visualization of Ppe1-mCherry and Sep3-GFP localizations at 11 hpi.

Video S3. 3D visualization of Ppe1-mCherry ring surrounding the Sep3-GFP small ring at 19 hpi.

Video S4. 3D visualization of Ppe1-mCherry ring surrounding the penetration peg at 22 hpi.

Video S5. 3D visualization of Ppe1-mCherry ring and papilla callose ring at the penetration site at 22 hpi.

**Supplementary Table S1.**
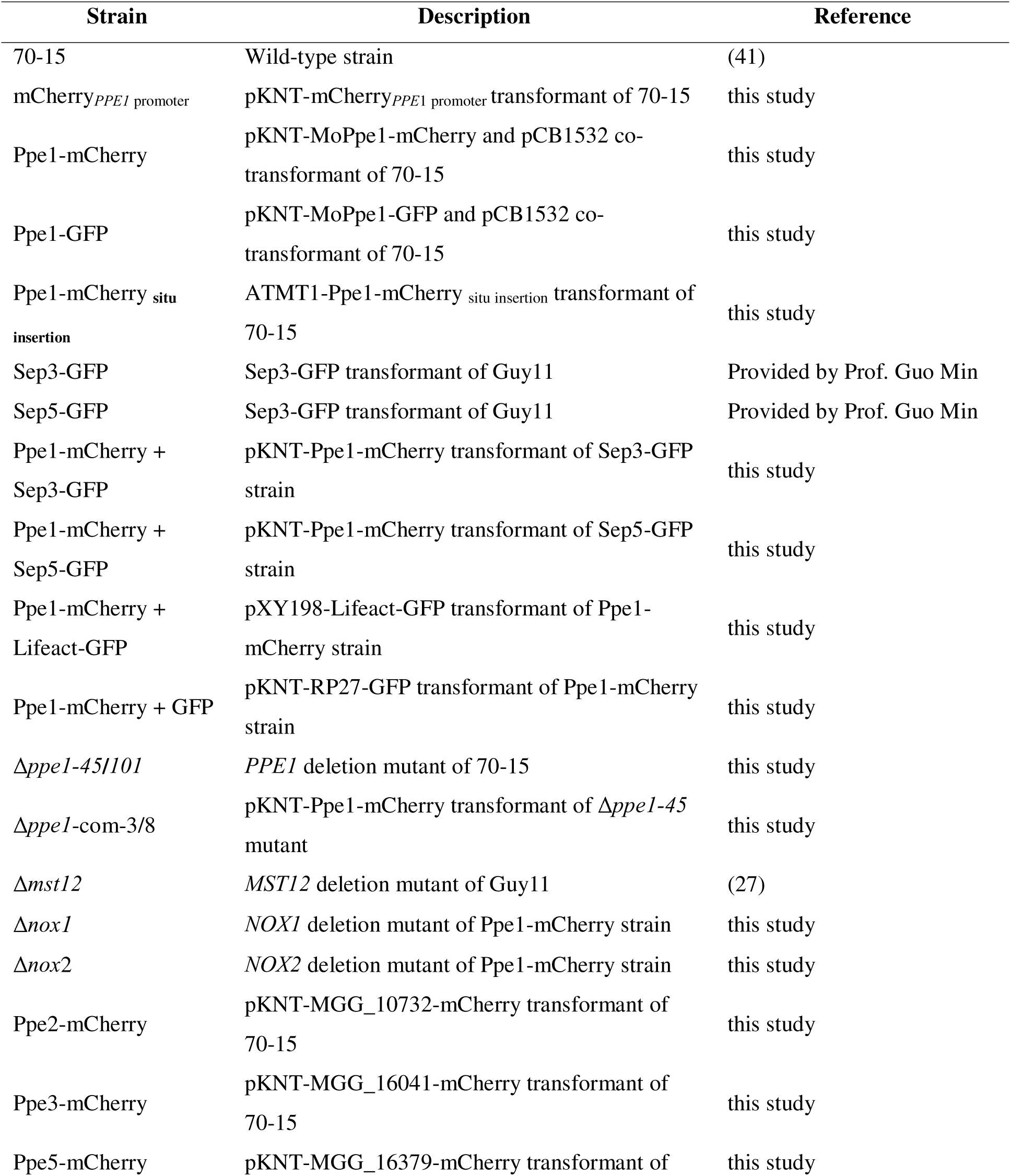
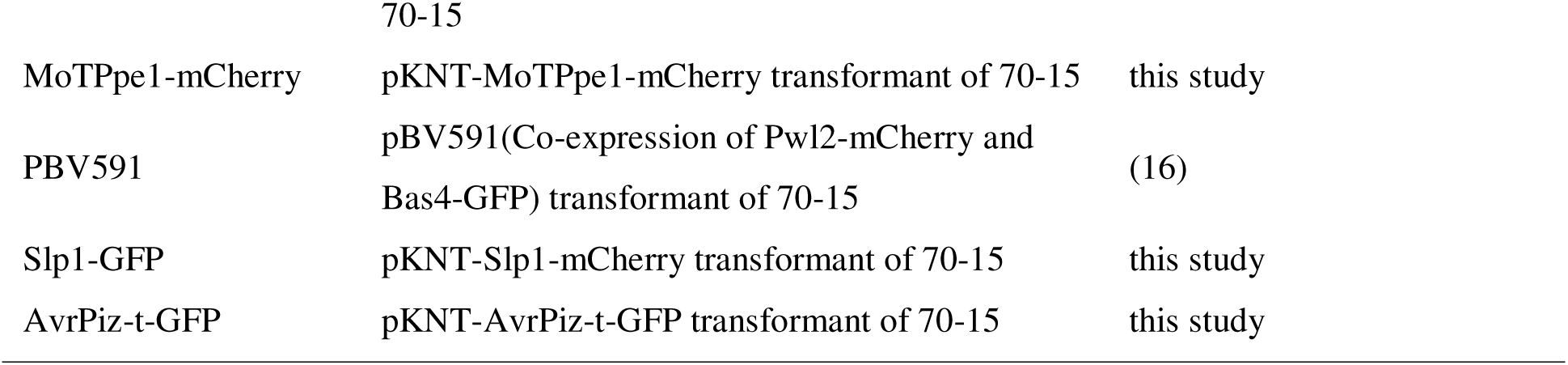
Fungal strains used in this study.

**Supplementary Table S2.**
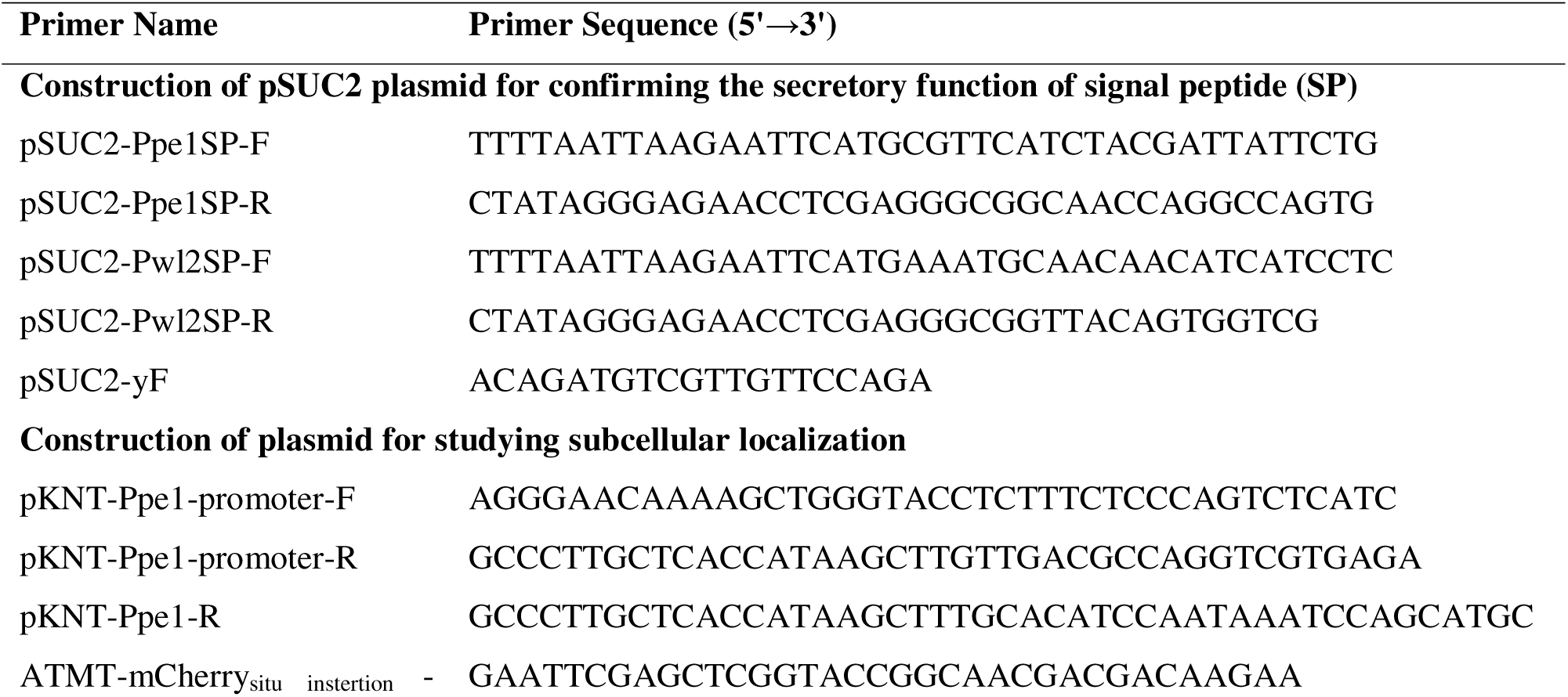
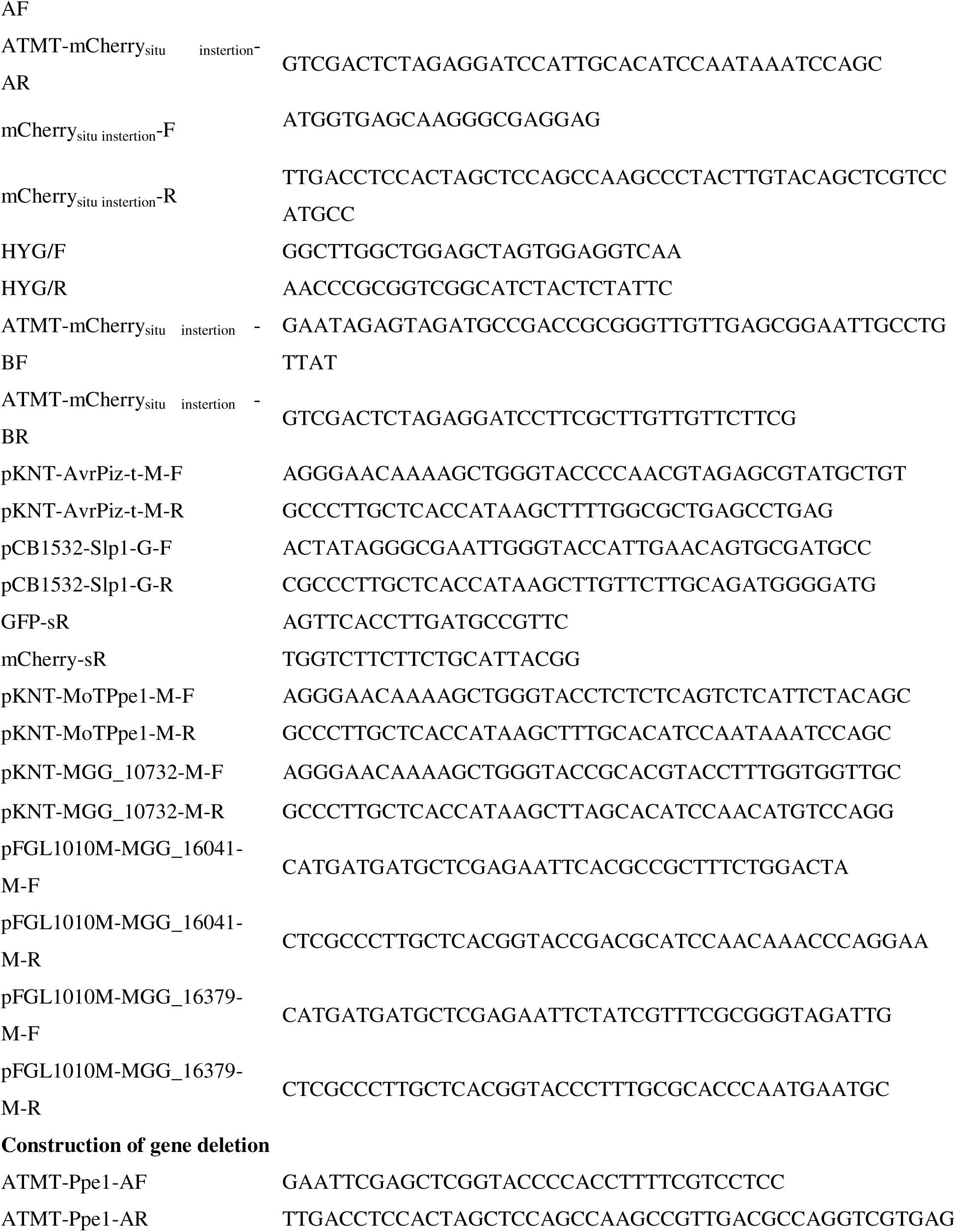
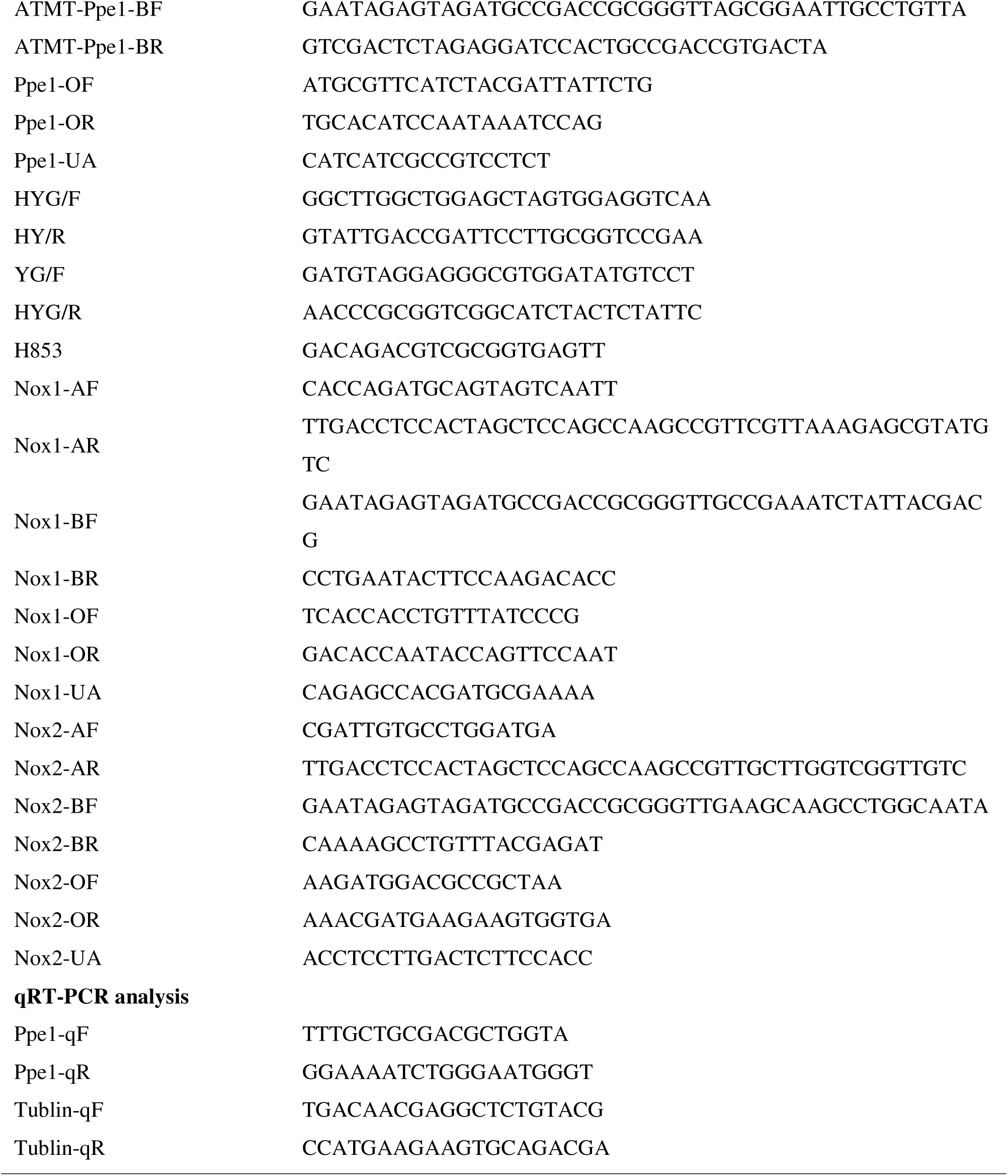
List of Primers used in this study.

